# A Sensory Memory to Preserve Visual Representations Across Eye Movements

**DOI:** 10.1101/2020.02.28.970277

**Authors:** Amir Akbarian, Kaiser Niknam, Kelsey Clark, Behrad Noudoost, Neda Nategh

## Abstract

During eye movements, the continuous flow of visual information is frequently disrupted due to abrupt changes of the retinal image, yet our perception of the visual world is uninterrupted. In order to identify the neuronal response components necessary for the integration of perception across eye movements, we developed a computational model to trace the changes in the visuospatial sensitivity of neurons in the extrastriate cortex of macaque monkeys with high temporal precision. Employing the model, we examined the perceptual implications of these changes and found that by maintaining a memory of the visual scene, extrastriate neurons produce an uninterrupted representation of the visual world. These results reveal how our brain exploits available information to maintain the sense of vision in the absence of visual information.

## Introduction

The question of how visual information acquired in a snapshot of the scene, during fixation, is integrated with the next snapshot after the eye moves, has puzzled psychophysicists, physiologists, and cognitive neuroscientists for decades (Binda and Morrone, 2018; Ibbotson and Krekelberg, 2011; Melcher and Colby, 2008; Wurtz, 2008). In order to understand the neural basis of visuospatial integration across saccadic eye movements (saccades), we recorded the spiking activity of 291 V4 and 332 MT neurons while the monkey performed a visually guided saccade task (Figure 1A, left, see STAR Methods and Figure S1). The animal maintained their gaze at a central fixation point (FP1) for 600-750 ms and upon the FP1 offset, shifted their gaze to a peripheral target (FP2) and fixated there for another 600-900 ms. Prior to, during, and after saccades, 7-ms small visual stimuli (probes) were presented pseudo-randomly within a matrix of 9×9 locations covering the FP1, FP2, and the estimated receptive fields of the neuron before and after the saccade (RF1 and RF2). In order to assess how extrastriate neurons represent the visual world we needed to trace the dynamics of their sensitivity as it changes during saccades. The neuron’s sensitivity (*g*) at a certain time relative to the saccade (*t*) is defined as the efficacy of a stimulus at a certain location (*x* and *y*) presented at a specific delay (*τ*) before that time to evoke a response in that neuron (Figure 1A, right). In order to assess this sensitivity map, we employed a computational approach. First, we decomposed the time and location into discrete bins of ∼3-6 degrees of visual angle (dva) and 7 ms time bins (resolution of probes). For the duration and precision of our experimental paradigm, a full description of a neurons’ sensitivity required evaluation of 10^7^ of these spatiotemporal units (STUs). We used a dimensionality reduction algorithm to select only those STUs that contribute to the stimulus-response correspondence (see STAR Methods; Figure S2). This unbiased approach excluded more than 99% of STUs, making it feasible to evaluate the contribution of the remaining ∼10^4^ STUs to the response of the neuron (8899.17±113.97 STUs per neuron). We developed a computational model to predict the neuron’s response based on an estimated sensitivity map. Using a gradient descent algorithm, we asked the model to determine the contribution (weight) of each STU of the sensitivity map, with the goal of maximizing the similarity between the model’s predicted response and the actual neuronal response (see STAR Methods). Figure 1B shows examples of weighted STUs for a model of an example MT neuron at a location inside its RF1 (top) and RF2 (bottom) for various times relative to the saccade. Note that the combination of STU weights across delays for a certain time and location is a representation of the neuron’s sensitivity, classically known as its “kernel” (middle panel). Overall, the model performed well in capturing the dynamics of neuronal responses, as well as providing high temporal resolution sensitivity maps of neurons (Niknam et al., 2019) (see STAR Methods; Figure S3-4).

**Figure 1.**
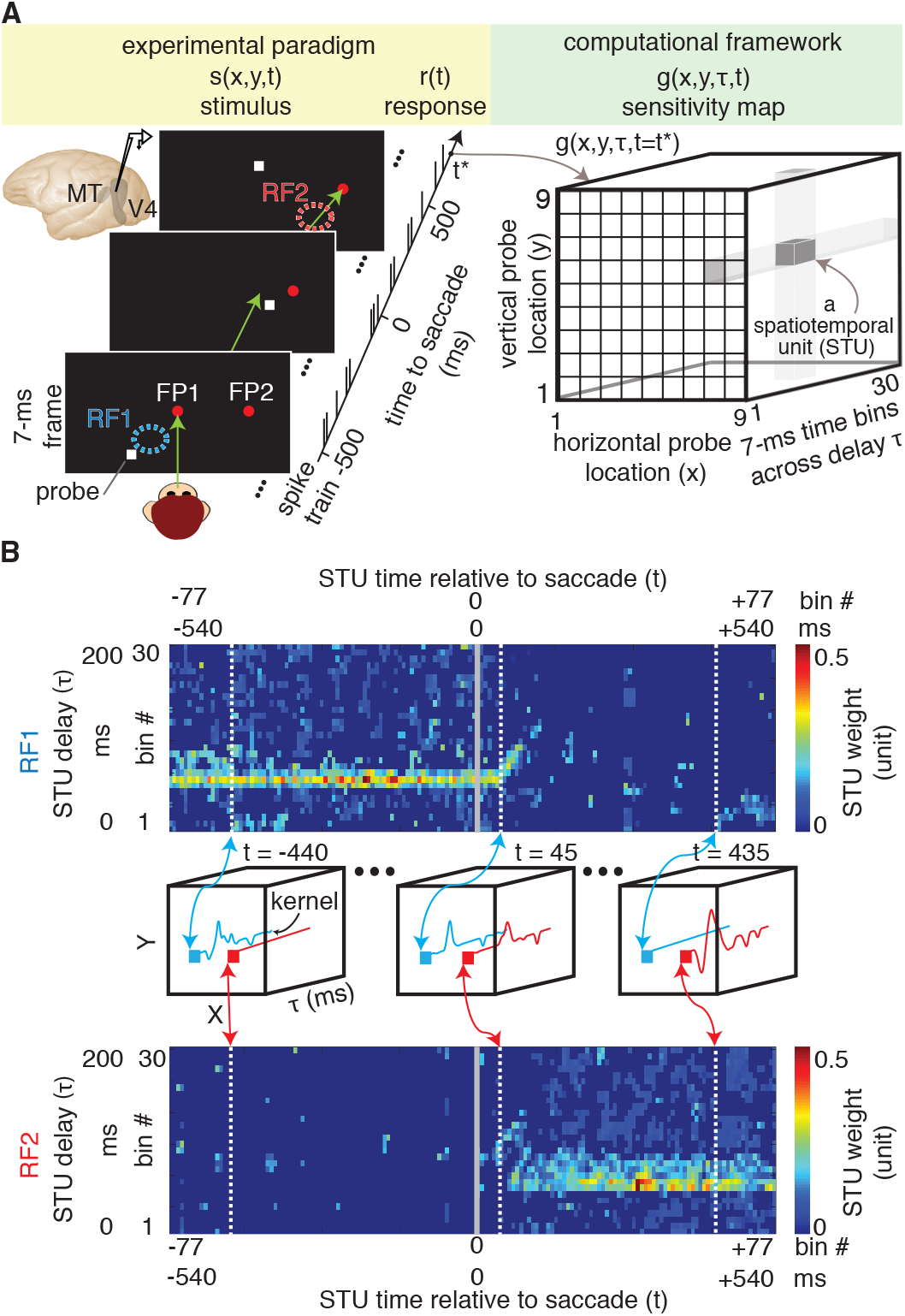
Decomposing sensitivity maps of neurons across saccades. (A) Experimental paradigm: monkey saccades from a fixation point (FP1) to a peripheral target (FP2). Throughout, visual probes (white squares) appear at pseudo-random locations. Green arrow shows gaze direction. Computational framework: neuron’s sensitivity map across delays and locations, illustrated schematically for one timepoint relative to saccade. (B) A sample neuron’s STU weights are shown for RF1 and RF2, across time to saccade and delay. The middle row shows example kernels (estimated as a weighted combination of STUs for RF1, blue, and RF2, red, for example pre-, peri- and post-saccadic timepoints).

The goal of our combined electrophysiology and computational approach is to identify the neural components underlying the integrity of perception across saccades. This requires translating the sensitivity map to a readout of the visual scene (employing the decoding aspect of the model), and then using this readout to assess the perceptual integrity around the time of saccades. By capturing the essential computations of the neuron, the model can be used to generate predictions about any unseen sequence of visual stimuli. An example of how this can be used to generate a readout of the visual scene is shown in Figure 2A. The model is used to predict responses to 9 probes around RF1 at various times relative to saccade onset. For a pair of probes, the spatial discrimination was then measured using the area under the curve (AUC) in the Receiver Operating Characteristic method based on the model-predicted responses (see STAR Methods; Figure S5A-D). Location discriminability is assessed by the average AUC across all pairs of probes, and is plotted for a single neuron at various times of its response (x-axis) for probes presented at different times relative to saccade onset (y-axis). Figure 2b shows the same location discriminability map for the population of 623 modeled neurons, and the blue contour indicates the times at which the response can differentiate between probe locations above a certain threshold (AUC>0.55). The same contour is shown in Figure 2c-top, along with the contour assessed with a similar method based on the location discriminability around RF2. Consistent with the subjective perception of a continuous visual scene, there is no time at which spatial sensitivity is lost during saccades, as indicated in the contact between the red and blue regions, and the overlap in their projections along the response time dimension (Figure 2c-top, overlap=23.23±4.92ms; see STAR Methods). The same phenomenon was also observed when assessing the detection performance of neurons (Figure 2c-bottom, see STAR Methods). Therefore, tracing the capacity of neurons to decode location information, the model predicts no gap between encoding information from the presaccadic scene and the postsaccadic one.

**Figure 2.**
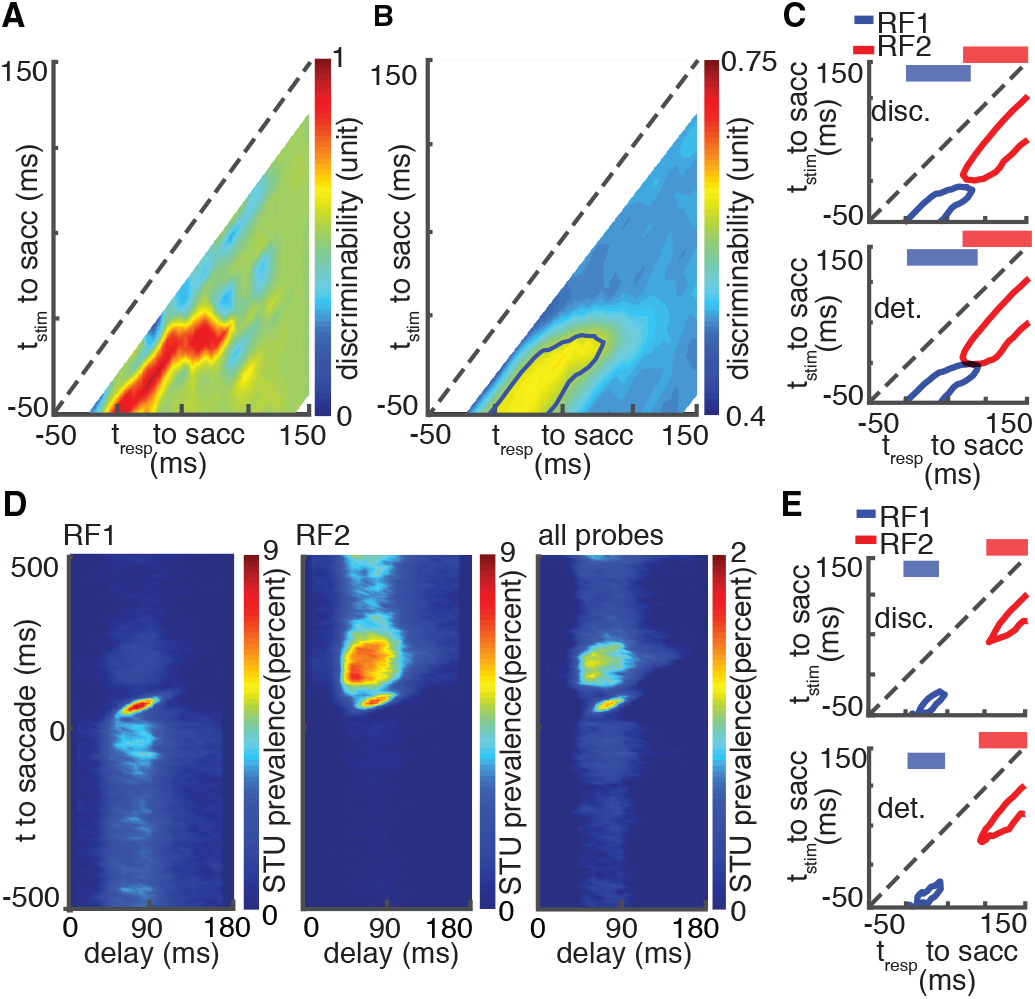
Quantifying the spatial integration and assessing the contributing perisaccadic modulatory components. (A) Location discriminability measured via model decoding for a sample MT neuron around its RF1. (B) Location discriminability around RF1 for the population of 623 neurons. (C) Temporal profiles of discriminability (top) and detectability (bottom) for RF1 and RF2; continuous sensitivity is reflected in the overlap in the projections on top. Contours indicate the times at which discriminability/detectability were above a certain threshold. (D) Temporal characteristics of modulated STUs. (E) Temporal profiles of discriminability (top) and detectability (bottom) for RF1 and RF2 after removing modulated STUs.

## RESULTS

The approach revealed an important insight about what exactly happens for the visual scene representation around the time of saccade. As shown in Figure 2b and c, responses up to 50 ms after saccade onset show that neurons consistently kept their spatial sensitivity to stimuli as early as 50 ms before that certain time (deviation from the line of unity, which is also a reflection of neuronal response latency. Interestingly, the blue curve is slightly farther from the line of unity around the time of the saccade (∼74ms deviation for response times of 50-90 ms), implying that the neuron loses its sensitivity to more recent stimuli and instead remains sensitive to stimuli presented earlier in time, a phenomenon which could contribute to filling the perceptual gap during saccades (see STAR Methods and Figure S5E-F for a verification at the neuronal level).

Having confirmed that the readout of the visual scene is indeed integrated across saccades and with a clue that this integration likely occurs by a change in the temporal dynamics of neuronal response we started our search to identify the exact extrastriate mechanism underlying this phenomenon. Importantly, the model provides the ability to independently manipulate individual components of neuronal sensitivity and assess their neuronal response consequences as well as their implication for visuospatial representation. This ability proved to be a very powerful tool in our quest. First, we identified the times at which saccades alter extrastriate neurons’ sensitivity by identifying the STUs whose contribution changes during saccades compared to fixation (see STAR Methods). The temporal distribution of saccade-modulated STUs is shown in Figure 2d for RF1, RF2, and all locations, across all modeled neurons. On average ∼26% of STUs were modulated during saccades (2342.89±45.36). Nulling these modulated STUs in the model, i.e. replacing their weights with the fixation weights, resulted in a clear gap in neurons’ sensitivity to visual information. Unlike the intact model in Figure 2c, the model lacking the perisaccadic modulations (Figure 2E) not only did not show any overlap between RF1 and RF2 sensitivity, it even implied a gap in the visuospatial representation by showing that for certain times, extrastriate neurons are not sensitive to either location (overlap=−36.88±6.18ms; see STAR Methods and Figure S6B). These results demonstrate the necessity of perisaccadic extrastriate changes for maintaining an integrated representation of visual space across saccades.

Numerous psychophysical phenomena happen during saccades: targets of eye movements are processed better, sensitivity to detect changes and displacements of other objects are reduced, and perception of time and space alters (Hamker et al., 2011). Although linked to several of these perceptual phenomena, and maybe even the ultimate goal of the system, maintaining an integrated representation of space is only one of the perisaccadic perceptual phenomena. Therefore, whereas Figure 2 verifies the necessity of perisaccadic sensitivity changes for this integration, we still need to determine which changes are specifically related to visuospatial integration. We defined the integration based on a model readout and then induce assumption-free alterations into the model to determine which of the modulated STUs (Figure 2D) are essential for an integrated representation of space, i.e. altering the model readout from Figure 2E to Figure 2C (See STAR Methods; Figure S6C). This unbiased search within the space of STUs revealed the times, delays, and locations of “integration-relevant STUs” (Figure 3A) (17.04±0.23% of the modulated STUs were integration relevant). More importantly, the model can then be used to link integration-relevant STUs to specific components of the neural response. For example, the black contours in Figure 3A highlight the integration-relevant STUs for RF1 and RF2 locations for all modeled neurons, and the black contours in Figure 3B highlight the stimulus-aligned response component generated by those specific integration-relevant STUs at RF1 and RF2. This approach isolated an alteration in the dynamics of responses to RF1 and RF2 stimuli presented within 10 ms of saccades as the neural substrate for an integrated representation of visual space. As indicated in the right panels, around the time of the saccade, the early part of the response to RF1 probes gradually disappears and a late response component emerges instead (which disappears after the eye has landed on the second fixation point). For RF2 probes a late component emerges first and gradually earlier components add to it to form the stimulus-aligned response of the neuron during the second fixation. The same phenomena observed in the model were also seen in the response of the population of neurons (Figure 3C). The phenomenon occurring at RF2 is reminiscent of the previously reported future field remapping (see STAR Methods; Figure S7) (Colby and Goldberg, 1992; Nakamura and Colby, 2002). The elongation of the RF1 response, which we call ‘RF persistence’, is however a novel finding and provides reassurance that our unbiased search is casting a wide net to identify the neural basis for an integrated representation of visual space.

**Figure 3.**
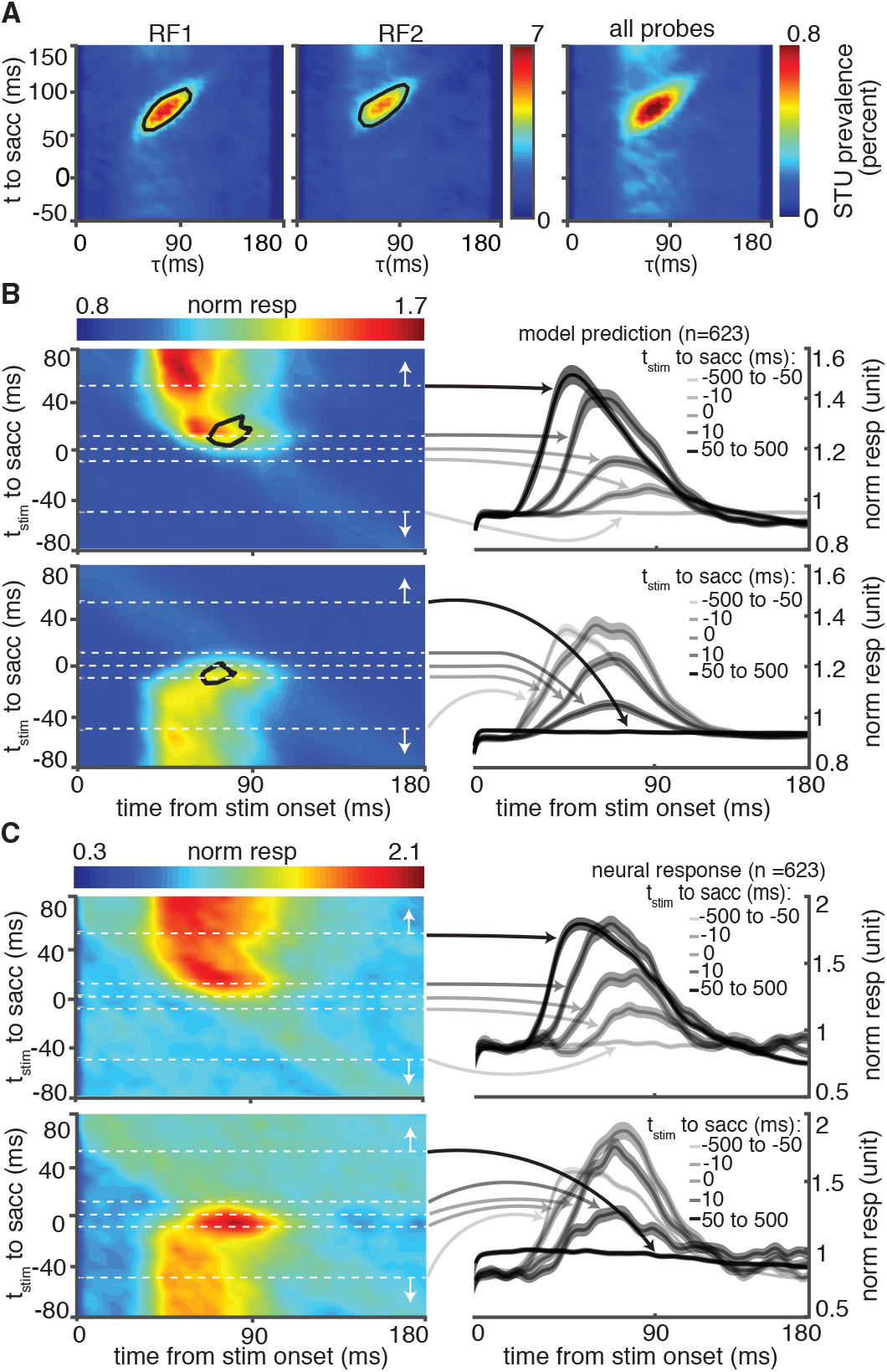
Identifying the neural basis for an integrated representation of visual space. (A) Temporal characteristics of integration relevant STUs. (B) Model-predicted response of the population across time to saccade and delay. The black contour indicates the response driven by integration-relevant STUs. PSTH traces on the right show the average population model-predicted firing rate, at five times relative to the saccade (dashed white lines on the left). (C) Plots are as in B, but with actual neuronal responses.

The phenomenon of RF persistence was observed in populations of both V4 and MT neurons. Figure 4a shows the rastergram and the average response of sample MT (left) and V4 (right) neurons. The average response of these neurons during 75-105 ms after probe onset increased by a factor of 6.37 and 2.31 for stimuli appearing around the saccade compared to fixation (MT_fixation_=13.85±0.91, MT_saccade_=88.28±9.78 sp/s, p<0.001; V4_fixation_=25.09±1.04, V4_saccade_=58.06±8.13 sp/s, p<0.001 Wilcoxon rank-sum test). As shown in Figure 4B, C both V4 and MT populations exhibited an enhanced late response to their RF stimulus around the saccade compared to fixation (MT_fixation_=40.97±1.47, MT_saccade_=45.39±1.71, p<0.001; V4_fixation_=36.15±1.61, V4_saccade_=45.66±1.89, p<0.001). This enhanced late response was accompanied by a suppression of early responses in both V4 and MT, although the precise timing of this phenomenon was slightly different between the two areas (see STAR Methods; Figure S8). In order to examine the spatial selectivity of the observed phenomenon, for each probe we measured the perisaccadic modulation index (PMI) as the difference between saccade and fixation responses (75-105ms after the stimulus) divided by their sum. PMI for RF probes was significantly greater than PMI for control probes outside the RF (ΔPMI_v4_=0.04±0.01, p=0.005; ΔPMI_MT_=0.04±0.01, p<0.001; Figure 4D). We also confirmed that the RF persistence phenomenon in MT is independent of whether the saccade direction is congruent or incongruent with the preferred motion direction of the neuron, ruling out saccade-induced retinal motion as the source of this phenomenon (PMI_congruent_=0.01±0.01, p=0.025; PMI_incongruent_=0.03±0.02, p=0.031, p_congruent vs. incongruent_ = 0.64, Wilcoxon rank-sum test; Figure 4E; See STAR Methods and Figure S9).

**Figure 4.**
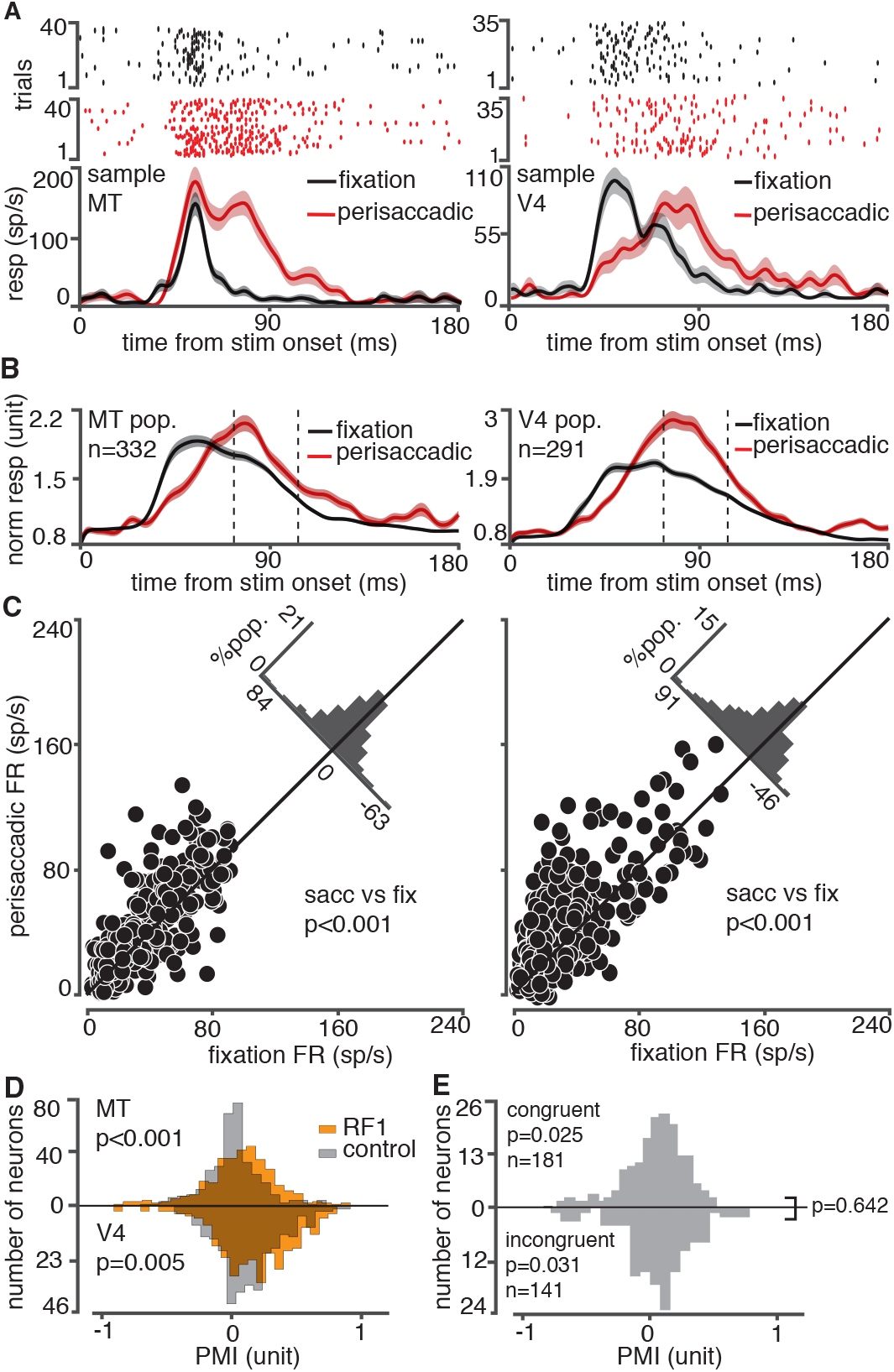
Late modulation of perisaccadic RF1 response. (A-B) Responses of sample neurons (A) and the neural population (B) to perisaccadic versus fixation RF1 probes in area MT (left) and V4 (right). Dashed lines indicate the late response window used in c-e. (C) The average late response of MT (left) and V4 (right) neurons to perisaccadic versus fixation RF1 probes. (D) PMI values for RF1 (orange) and control probes (gray), for the MT (top) and V4 (bottom) populations. (E) PMI values for saccade directions congruent (top) versus incongruent (bottom) with MT neurons’ preferred direction. pop., population.

## DISCUSSION

Combining physiological and modeling approaches, we developed a tool to assess how the visual scene is encoded in the responses of extrastriate neurons. We found that these neurons are capable of ‘stitching’ their presaccadic representation to the postsaccadic one by maintaining a memory of the scene. Prior to saccade, response of extrastriate neurons at a certain time represents visuospatial phenomena occurring ∼50 ms before. When the eye moves and the flow of retinal information is disrupted, a gap occurs in visuospatial representation and instead of representing the visual events 50ms ago, extrastriate neurons maintain a representation of events further back in time (∼75ms). Therefore, extrastriate neurons do go through a representational gap but are able to fill in the gap and provide an integrated representation. The computational approach not only revealed this compensatory mechanism, it also enabled us to target the exact neuronal response changes essential for this compensation by identifying the phenomenon of RF persistence, a sensory memory mechanism which can preserve information across saccades.

This discovery of RF persistence dovetails with previous psychophysical studies suggesting the necessity of such memory mechanism to preserve vision throughout the brief periods that the visual signal is lost during saccades or blinks (Coltheart, 1980; Rensink, 2014). Importantly, psychophysics experiments have implied that the preservation of vision across saccades might rely on mid to high-level visual areas rather than earlier parts of the visual hierarchy (Burr and Morrone, 2005; Melcher, 2005). Moreover, observing the RF persistence phenomenon in both V4 and MT implies that this sensory memory is a characteristic of the visual system independent of whether the signal originates from chromatic/achromatic or motion sensitive pathways earlier in the visual stream (Merigan and Maunsell, 1993). However, how exactly an extrastriate neuron knows when to elongate its response remains unknown. The intrinsic signal within these area (e.g. due to abrupt change in flow of visual information) might be enough but these areas also receive a copy of motor command (e.g. via the tectopulvinar pathway to MT(Lyon et al., 2010)) as well as attentional signal (via direct projections from the Frontal Eye Field (Merrikhi et al., 2017; Noudoost and Moore, 2011). Considering that V4 is expected to receive the motor command through MT and the dynamics of response changes in MT did not lead those in V4 (STAR Methods; Figure S8C), the role of top-down and intrinsic signals and their interactions need to be a focus of future studies.

It is imperative to emphasize that the phenomenon of perceptual stability, the subjective experience of a stable world during saccades, might require more than an integrated sensory representation. The stability has been shown to rely on working memory mechanisms (Irwin and Andrews, 1996; Peterson et al., 2001), and information outside a retinotopic framework might also be involved (Zirnsak et al., 2014) (see STAR Methods; Figure S10). The current results, however, show clearly that for a short period of time, retinotopic visual areas are able of reconstructing the visual world using available information, a resource that can be employed by other areas and frames of reference to generate a stable, uninterrupted sense of vision.

## AUTHOR CONTRIBUTIONS

N.N and B.N conceived the study. B.N. performed the surgical procedures. B.N., N.N. and A.A. designed the experiment. B.N. and A.A. performed the physiology experiments and acquired data. A.A. analyzed the neural data. A.A., K.N. and N.N. designed, developed and validated the computational model. B.N., N.N., A.A. and K.C. wrote the manuscript.

## ACKNOWLEDGMENTS

We would like to thank the animal care personnel at the University of Utah. We specifically thank Rochelle D. Moore and Dr. Tyler Davis for their assistance in performing the experiments. Funding: This work was supported by the startup fund from Montana State University and University of Utah to NN and BN; NIH R01EY026924 and NSF1439221 to BN and NSF1811543 to NN and an Unrestricted Grant from Research to Prevent Blindness, Inc. to Moran Eye Center.

## STAR⋆METHODS

## LEAD CONTACT AND MATERIALS AVAILABITIY

Further information and requests for resources should be directed to and will be fulfilled by the Lead Contacts, Neda Nategh and Behrad Noudoost. This study did not generate new reagents.

## EXPERIMENTAL MODEL AND SUBJECT DETAILS

### Animal Preparation

Four adult male rhesus monkeys (monkeys B, P, E, and O; Macaca mulatta) were used in this study. All experimental procedures complied with the National Institutes of Health Guide for the Care and Use of Laboratory Animals and the Society for Neuroscience Guidelines and Policies. The protocols for all experimental, surgical, and behavioral procedures were approved by Institutional Animal Care and Use Committees of the University of Utah and Montana State University. Animals were pair-housed when possible and had daily access to enrichment activities. During the recording days, they had controlled access to fluids, but food was available ad libitum.

### Surgical procedures

For each animal, a head-post was implanted on the skull using dental acrylic and orthopedic titanium screws. The surgical procedures were performed under strict aseptic conditions while the animals were anesthetized by Isoflurane. Recording chambers were mounted on the skull and fastened by screws and dental acrylic. The craniotomy was performed within the chamber, giving access to extrastriate visual areas including V4 and MT.

### Experimental paradigm and data acquisition

Monkeys performed a visually guided saccade task during which task-irrelevant pseudorandom square stimuli flashed on the screen (Figure 1). Four adult male rhesus monkeys were trained to fixate on a fixation point (FP1; a central red dot) located in the center of the screen. After they fixated, a second target (FP2; a peripheral red dot) appeared 10-15 degrees away. Then, after a randomized time interval between 600 and 750 ms (drawn from a uniform distribution), the fixation point disappeared, cuing the monkeys to make a saccade to FP2. After remaining fixated on the FP2 for 600 to 900 ms monkeys received a reward. During this procedure, a series of pseudo randomly located probe stimuli were presented on the screen in a 9 by 9 grid of possible locations. Each stimulus was a white square (full contrast), 0.5 by 0.5 degrees of visual angle (dva), against a black background. Each stimulus lasted for 7 ms and stimuli were presented consecutively without any overlap, such that at each time point there was only one stimulus on the screen. The locations of consecutive probe stimuli followed a pseudorandom order, called a condition. In each condition, a complete sequence of 81 probe stimuli was presented throughout the length of a trial. Conditions were designed to ensure that each probe location occurred at each time in the sequence with equal frequency across trials. For each recording session, the grid of the possible locations of the probes was positioned such that it covered the estimated pre- and post-saccadic receptive fields (RFs) of the neurons under study, as well as the FP1 and FP2. The spatial extent of the probe grids varied from 24 to 48.79 (mean ± SD = 40.63 ± 5.93) dva horizontally, and from 16 to 48.79 (mean ± SD = 39.78 ± 7.81) dva vertically. The (center-to-center) distance between two adjacent probe locations varied from 3 to 6.1 (mean ± SD = 5.07 ± 0.74) dva horizontally, and from 2 to 6.1 (mean ± SD = 4.97 ± 0.97) dva vertically. For the MT neurons, the motion direction preference was assessed using a full field Gabor paradigm before the saccade task. The monkey maintained fixation while a full field Gabor stimulus, moving in one of 8 directions, was displayed for 800 ms.

Throughout the entire course of the experiment, the spiking activity of the neurons in areas V4 and MT was recorded using a 16-channel linear array electrode (V-probe, Plexon Inc., Dallas, TX) at a sampling rate of 32 KHz, and sorted offline using the Plexon offline spike sorter and Blackrock Offline Spike Sorter (BOSS) software. The spike sorter program was employed to perform a principal component analysis, and clusters of spikes with similar waveform properties were manually classified as belonging to a single neuron (single unit). The sorted spikes were then read into Matlab to verify the presence of a visually-sensitive RF. From a population of 709 well-isolated neurons, 86 neurons were discarded because they did not respond to any probe stimuli before and/or after the saccade, and the rest were used for analyses. The eye position of the monkeys was monitored with an infrared optical eye tracking system (EyeLink 1000 Plus Eye Tracker, SR Research Ltd., Ottawa, CA) with a resolution of < 0.01 dva, and a sampling frequency of 2 KHz. Stimulus presentation in the experiment was controlled using the MonkeyLogic toolbox (Hwang et al., 2019). Visual stimuli were presented on a 24-inch ASUS VG248QE LED monitor with a resolution of 1920×1080 pixels with a refresh rate of 144 Hz, positioned 24-30 cm in front of the animal’s eyes. A photodiode (OSRAM Opto Semiconductors, Sunnyvale CA), mounted on the lower left corner of the monitor, was used to record the actual onset and offset times of stimuli appearing on the screen with a continuous signal sampled and stored at 32 KHz.

In total, data were recorded from 332 MT and 291 V4 neurons during 108 recording sessions. In 76 recording sessions, the saccades were made to the left (439 of 663 neurons), and in 23 recording sessions, the saccades were made to the right (146 of 663 neurons), and in the remaining 9 sessions (38 neurons) the saccades were made in other directions (195, 200, and 225 degrees). The positioning of the probe grids, the spatial distribution of the RF of the neurons, and the average of the photodiode signal for an example session is illustrated in Figure S1.

### RF estimation

The RF1 and RF2 locations used to calculate detectability and sensitivity in Figure 2 refer to the probe locations that generated the maximum firing rate during the fixation period before and after the saccade, respectively. For each probe location, the probe-aligned responses are calculated by averaging the spike trains over repetitions of the probe before or after the saccade (greater than 100 ms before or after the saccade onset), from 0-200 ms following probe presentation, across all trials. The response is then smoothed using a Gaussian window of 5 ms full width at half maximum (FWHM). For illustration of the distribution of RF locations on the screen (Figure S1C), the probe-aligned responses are averaged over a window of 25 ms around the time of maximum response for each neuron. The map is then interpolated (interleaving 1000 points between each probe location) and thresholded at 0.99 of the maximum response. The center of RF is then calculated as the center of mass of the resulting spatial map. For the population of neurons, the average RF center was located 8.35±0.16 dva away from the FP1 (Figure S1C).

## METHOD DETAILS

### Dimensionality reduction for computing neuron’s time-varying sensitivity map

The fast, complex dynamics of changes in the neurons’ spatial sensitivity across a saccadic eye movement demand a high-dimensional representation of neurons’ spatiotemporal kernels in order to capture those perisaccadic dynamics. For any time relative to saccade onset the set of stimuli driving the response can be described in terms of their location (*X* and *Y*) and the delay between the stimulus presentation and the response time (*τ*). The goal is to determine this sensitivity map and trace its changes across time (Figure 1B). In our experiment, for a 200 ms delay kernel across 1000 ms of response time, this space could be decomposed into ∼10^7^ spatiotemporal units (STUs; Figure 1C). Since the stimulus presentation resolution is 7 ms, we represent the variation of sensitivity across the time dimensions using a set of temporal basis functions, 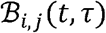, whose centers are separated by 7 ms across *τ* and *t* dimensions (Eq. 1). This way we down-sample the time into a sequence of binned STUs whose values can change every 7 ms.

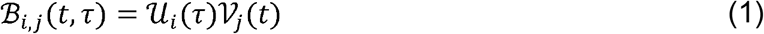

where 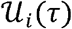 and 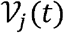 are chosen to be B-spline functions of order two. 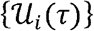 span over the delay variable *τ*, representing a 200 ms-long kernel using a set of 33 knots uniformly spaced at {−13, −6, …, 204, 211} ms (in total, 30 basis functions), and 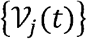 span over the time variable *t*, representing a 1081 ms-long kernel centered at the saccade onset using a set of 159 knots uniformly spaced at {−554, −547, …, 545, 552} ms (in total, 156 basis functions).

This representation reduces the dimensionality of the spatiotemporal sensitivity map by about two orders of magnitude, however, it is still far beyond the practical dimensionality for a computationally robust estimation of the sensitivity values (Niknam et al., 2019) using an experimentally tractable amount of data. The short duration of saccade execution makes it infeasible to acquire a large number of data points from all spatial locations and times relative to saccade onset. To address this, we use a statistical approach to identify the STUs whose presence significantly contributes to the neuron’s response generation at a given time. Figure S2 shows this pruning procedure. For each STU we compare the distribution of its weight estimated by fitting a generalized linear model (GLM) on 100 subsets of randomly chosen spike trains (35% of total trials) versus the control distribution obtained using 100 subsets of shuffled trials in which the stimulus-response relationship was distorted. The conditional intensity function (CIF) of this GLM is defined as,

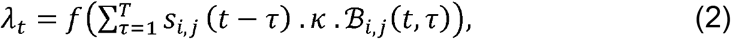

where *λ_t_* is the instantaneous firing rate of the neuron, *s_i,j_* is the stimulus history of length *T* at location (*i,j*), *κ* is the weight of a single STU, represented by basis function 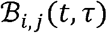, whose contribution significance is evaluated. An STU is discarded if the mean of these two weight distributions fails to satisfy the following condition:

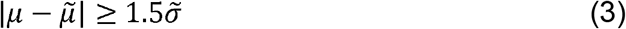

where 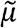 and 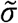 are respectively the mean and standard deviation of the control weights distribution, and *μ* is the mean of the original weights distribution. This pruning process reduces the dimensionality of STU space to ∼10^4^, which makes fitting of our encoding model to the sparse perisaccadic spiking data feasible and prevents an overfitted result. We then use only this subset of STUs to parameterize the linear filtering stage of an encoding model in a much lower dimensional space in order to determine the weights with which these STUs are combined to generate the neuron’s spatiotemporal sensitivity (Figure 1B); at each timepoint relative to the saccade, the weighted combination of these STUs over probe locations and delay times describes the neuron’s sensitivity kernels (Figure 1B, middle panel) defined as,

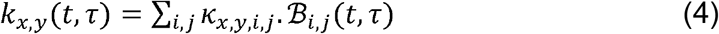

where {*κ_x,y,i,j_*} are the weights of the STUs obtained by estimating the encoding model (defined in Eq. 5 below). Note that the summation is over the subset of 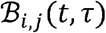 whose corresponding STU was evaluated significant according to in equation 3, while the weights for the remaining STUs were set to zero.

This low-dimensional set of the selected STUs enabled us to fit our encoding model to the sparse perisaccadic data and characterize the encoding principles of each neuron at each time relative to the saccade.

### Encoding model framework and estimation

#### Encoding model framework

In order to estimate the weight of each STU, we develop a new variation of classical GLMs, termed the sparse-variable GLM (S-model (Niknam et al., 2019), Figure S3), which enables us to represent a high-dimensional model using a sparse set of variables selected through a dimensionality reduction process and estimate those variables. Using this encoding model, we can capture the neuron’s high-resolution spatiotemporal sensitivity using limited perisaccadic data. Using dimensionality reduction to identify the subset of STUs representing the stimulus-response relationship is a key step enabling us to successfully fit the S-model parameters using the sparse neuronal data during saccades. Using this set of STUs we parameterize the stimulus kernels *k_x,y_*(*t,τ*) in the CIF of the S-model defined as,

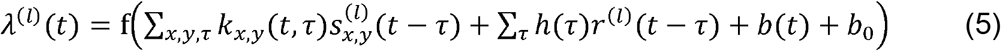

where *λ*^(*l*)^(*t*) represents the instantaneous firing rate of the neuron at time *t* in trial *l*, 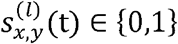 denotes a sequence of probe stimuli presented on the screen at probe location (*x,y*) in trial *l* with 0 and 1 representing respectively an off and on probe condition, *r*^(*l*)^(*t*) ∈ {0,1} indicates the spiking response of the neuron for that trial and time, *k_x,y_*(*t,τ*) represents the stimulus kernel at probe location (*x,y*), *h*(*τ*) is the post-spike kernel applied to the spike history which can capture the response refractoriness, *b*(*t*) is the offset kernel which represents the saccade-induced changes in the baseline activity, *b*_0_ = f^−1^(*r*_0_) with *r*_0_ defined as the measured mean firing rate (spikes per second) across all trials in the experimental session, and finally,

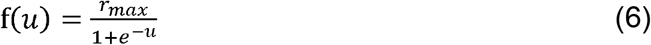

is a static sigmoidal function representing the response nonlinear properties where *r_max_* indicates the maximum firing rate of the neuron obtained empirically from the experimental data. This choice of the neuron’s nonlinearity is consistent with an empirical nonlinearity estimated nonparametrically from the data. All trials are saccade aligned, i.e., *t* = 0 refers to the time of saccade onset. Then using an optimization procedure in the point process maximum likelihood estimation framework, we fit the model to sparse spiking data at the level of single trials. The resulting encoding framework enables us to decipher the nature of saccade-induced modulatory computations in a precise and computationally tractable manner using the time-varying kernels representing the neuron’s dynamic sensitivity across different delays and locations for any specific time relative to the saccade.

#### Model fitting

We model the spiking response using a conditionally inhomogeneous Poisson point process, with its CIF defined in Eq. 5. The probability of a spike train associated with this Poisson process is given by,

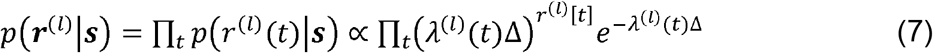

where ***s*** is the sequence of input stimuli, and ***r***^(*l*)^ = {*r*^(*l*)^(*t*)} represents the sequence of binned spike counts with bins of size Δ ms on trial *l*. Here, the bin size was chosen equal to 1 ms which ensures that at most one spike can fall in each time bin. The point process log-likelihood (*LL*) of the observed spike trains given the model is as follows,

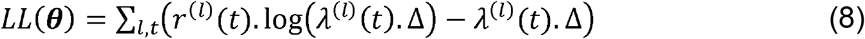

where ***θ*** denotes the set of parameters used to describe the model kernels defined in Eq. 5. Note that *κ_x,y,i,j_*, representing the weights of STUs in the stimulus kernels, is among this set of parameters. The parameters in ***θ*** are estimated by maximizing the log-likelihood function in Eq. 8. The sigmoidal nonlinearity in the S-model’s CIF (Eq. 5) makes the *LL* function not convex, meaning it may not give a unique optimal solution (more details in (Akbarian et al., 2017; Niknam et al., 2019)). Also, considering the number of data points (spiking events) relative to the number of model parameters to be estimated, this optimization may be subject to the overfitting problem. To handle these nonconvexity and overfitting challenges and find a robust and interpretable solution among other possible solutions, we adopted several parameter selection, sparsity, and smoothness regularization strategies. These strategies included early stopping, representation of model kernels using smooth basis functions, data-informed initialization, an iterative, robust gradient-based search algorithm, and cross validation, in addition to those strategies described in the dimensionality reduction section for the stimulus kernels specifically. For more details of these optimization strategies refer to (Niknam et al., 2019).

#### Model evaluation

The models’ performance is evaluated over test data, which is used neither for training the model nor for the validation process. In order to estimate the instantaneous firing rates, the sequences of stimuli presented to the neuron are given to the model according to Eq. 5, and the spiking history input to the post-spike kernel is simulated, similar to (Niknam et al., 2019). The performance of the model in predicting the neural response is assessed at the level of both firing rate prediction (Figure S3A-C) and also single trial, single spike prediction (Figure S3D-E).

Figure S4A shows how well the model captures the stimulus-response relationship by evaluating its ability to capture the average firing rate in response to the repeated presentation of a probe in the neuron’s RF during the fixation period (500 to 200 ms before saccade) for a sample MT neuron. We then quantify the similarity between actual and model-predicted responses across the population of recorded neurons using the explained variance (EV) measure (Schoppe et al., 2016). Figure S4B shows that the EV between the model-predicted firing rate and the empirical firing rate (y-axis) matches that obtained between 1000 pairs of average firing rate sequences measured over randomly selected subset of probe-aligned spike trains, used as a measure of inherent variability in the neural data itself (x-axis). The average firing rate sequences were computed by binning the probe-aligned spike response using nonoverlapping windows of 30 ms and smoothing the binned response with a Gaussian window of 5 FWHM and normalizing to have a mean of zero and unit standard deviation. The model-data EV and data-data EV are highly correlated, showing that the model-predicted response captured the stimulus-response relationship in the data across the population of the neurons (data-data EV: 84.79±0.54, model-data EV: 79.33 ± 0.78, Pearson correlation: 0.85, p<0.001).

Next, we generalized this firing rate level accuracy analysis by evaluating how well the model predicted the firing rate in response to the presentation of experimental sequences of probe stimuli appearing at random locations during the fixation period (500 to 200 ms before saccade). Figure S4C shows that the correlation coefficient (CC) between the model-predicted firing rate and the empirical firing rate in response to the repeated presentation of a sequence of probe stimuli falls within the level of the inherent trial-by-trial variability. The data-data correlation coefficient was measured between binned firing rates in response to the same 300 ms stimulus sequence; data were randomly split (60%-40%) 15 times and the mean is reported. The binning, smoothing, and normalizing of these data were the same as in the EV analysis. The normalized correlation in percent is calculated as the ratio between the model-data correlation (y-axis) and the data-data correlation (x-axis) and is shown as the diagonal histogram in Figure S4C. The data-data and model-data CC are positively correlated (data-data CC= 0.46± 0.005 and for the model-data CC it is 0.32± 0.006, Pearson correlation= 0.91, p<0.001) showing that the model is able to capture the trial-by-trial variability in the data where exactly same sequence of stimuli were presented to the neuron.

The results show that the model follows the neural response at the single trial level as shown in Figure S4D. The best and average fit trials in Figure S4D are chosen based on the largest and median normalized log-likelihood (LL) of trials for each sample neuron. We also analyzed the performance of our model on predicting single spikes and single trials, by assessing how well the model-predicted firing rate matches the observed spiking data on individual trials. Figure S4E shows the model’s performance in predicting the neural response in the perisaccadic period (from 0 to 150 ms after the saccade onset) versus that measured during fixation (−300 to −150 ms relative to saccade onset) by comparing the normalized log-likelihood of the model prediction in these time periods. The normalized LL, calculated as the LL of the spike trains using the predicted firing rate under the model minus that under a null model and normalized by spike counts, as reported in (Akbarian et al., 2017; Niknam et al., 2019), indicates the amount of information being conveyed by individual spikes and evaluates how closely the model describes the timing of recorded spikes. To calculate the normalized LL, the null model is assumed to be a model where the instantaneous firing rate of the neuron is set to its average firing rate. The distribution of the normalized LL values across the population of neurons shows that the model performance in describing neural data in the perisaccadic period is as good as during the fixation period (perisaccadic normalized LL= 0.17±0.00 bits/sp, fixation normalized LL=0.17±0.00 bits/sp, p=0.75).

### Details of discriminability and detectability analysis

The decoding aspect of the model enables us to develop a readout of the visual scene using the model-predicted responses. The model readout provides a detailed description of the neural decoding capability across a saccadic eye movement, which can be used to trace a specific perceptual phenomenon (in our case, visuospatial integration across saccades) and test the specific components of the neural response that the phenomenon relies on. By capturing the essential computations of the neuron, the model can be used to generate predictions about arbitrary sequences of visual stimuli not present in the experimental data. We have used this aspect of the model to predict how the decoding capability of the neural response changes across a saccade, in terms of its ability to detect the presence of a particular probe. The detectability of an arbitrary probe is measured by evaluating the ability to detect the presence of that particular probe from the model-predicted response; i.e., when that probe is presented (ON) versus when it is not (OFF). The detectability of probes can differ based on the time between the probe presentation and the time of response that the decoding is being based on (referred to as the delay). At any time in the neural response (denoted as *t* in Figure S5A), the probe is only detectable if it is presented within a certain delay range (*τ*; we evaluated delay values from 0-200 ms). During the fixation period, long before the saccade onset, the RF1 probe’s detectability is maximum around the latency of the neuron. However, during the perisaccadic period, the neuron becomes sensitive to different probes and with different latencies. Figure S5A shows how the detectability of the RF1 probe of a sample neuron (RF1 probe: *s*\*) is computed at an arbitrary time (*t*\*) where the probe is presented at (*t*\* – *τ*\*). To measure the detectability of *s*\* at *t*\*, *τ*\* the AUC measure (Macmillan and Creelman, 2004) has been used to evaluate the difference between the distribution of responses evoked at *t*\* (*λ_t_*\*) by the presence of *s*\* at *t*\* − *τ*\* versus in the absence of *s*\*, each embedded within a 200 ms random sequence of other probes. The model’s predicted response at time *t*\* (denoted as *λ*_*t*\*_) is generated for 100 random sequences of probes, when specific probe *s*\* is ON and 100 random sequences in which that probe is OFF, *τ*\* before *t*\* (i.e., at time *t*\* − *τ*\*). The detectability is then measured as AUC of the evoked response (*λ*_*t*\*_) to the ON versus OFF trials (histograms shown in Figure S5A). To calculate the average detectability, mean AUC was calculated across 20 repetitions for each time and delay combination, each repetition over a randomly selected 80% of ON and OFF trials. Figure S5B shows the detectability at a sample time *t*\* = +10 ms relative to the saccade where the RF1 probe is presented 140 to 20 ms before saccade onset (−150 – *τ* < −30 *ms*); at this time, the RF1 probe is detectable only when it was presented around 55 ms before the response (normal latency of the neuron). To track detectability across times and delays, the detectability of probes is measured at different values of *t* relative to the saccade and *τ* relative to each response time (*t*: 50 ms before to 300 ms after saccade with 10 ms steps, and *τ*: 190 to 30 ms before response time with 10 ms steps, Figure S5C). The detectability map of the neuron for each probe location provides a quantification of the decoding capacity of the neuron across the eye movement (Figure S5C shows the map for the RF1 probe of a sample neuron); the shift in detectability from RF1 to RF2 is shown in Figure S5D. Over time, the probe location with the highest detectability shifts from RF1 to RF2, as shown for the population of neurons in Figure S5D; the contours show the times and delays at which one can detect the presence of the stimulus based on the response of the neuron, i.e., the AUC values are above a threshold of 0.61.

In a similar way, we used the decoding approach to measure the location discriminability of the neural response, in terms of ability to discriminate a probe from the immediately surrounding probes using the model-predicted response. To measure the location discriminability at each probe location, 100 random sequences were presented to the model with the center probe presented at a specific delay, and AUC was measured versus 100 trials where one of the adjacent probes was presented at the same delay. Mean sensitivity for each of the surrounding probes was then calculated across 20 AUC measurements, each using 80% of trials. The location discriminability reported in Figure 2A, B, C and E are then the average of discriminability over 8 surrounding probes around the RF1 or RF2 probe. The thresholds used in Figure 2B, C, and E are ROC > 0.57 for discriminability and ROC>0.61 for detectability.

We also calculated the detectability and discriminability maps based on the recorded neural responses. Figure S5E-F illustrates the detectability and discriminability maps calculated based on the recorded neural responses and averaged over the population of neurons. To calculate the detectability of a probe at an arbitrary time relative to the saccade (*t*) and at a specific delay (*τ*), the spike count of the neuron is averaged over a 10 ms window around time *t*, in the trials that the probe was presented in a 10 ms window around time *t* − *τ*. The detectability is then measured as the AUC of the ROC between the spike count in the trials where the probe was presented versus trials where it was not. Similarly, for the discriminability, we evaluate the ability of the neuron’s response at time *t* to differentiate a probe from its surrounding probes in terms of the spike count of the neuron in a 10 ms window around time *t* in the trials that probes were presented at in a 10 ms window around *t* − *τ*.

### Identifying modulated and integration-relevant STUs

As discussed previously, only the STUs at specific times and delays are contributing in the neural response generation (green STU in Figure S2A). When the spatial and temporal sensitivity of a neuron changes during the perisaccadic period, the distribution of STUs (across times and delays) will be altered. We defined modulated STUs as those for which the prevalence of STUs in a 3×3 window around that STU’s time and delay is significantly different for stimuli presented perisaccadically vs. during fixation. Each STU is considered modulated if the prevalence of STUs fulfills following condition:

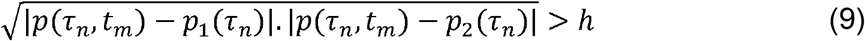

where *p*(*τ_n_,t_m_*) is prevalence of STUs in a 3×3 window around the *n*th bin of delay and mth bin in time 1 < *n* < 30,1 < *m* < 156, *p*_1_(*τ_n_*) is the prevalence of the STUs across fixation period before saccade calculated over bins 1 to 60 spanning 540 to 120 ms before saccade onset at nth bin of delay, 1 < *n* < 30, and *p*_2_(*τ_n_*) is the prevalence of STUs in the fixation period after saccade calculated over bins 120 to 156 spanning 280 to 540 ms after saccade onset at *n*th bin of delay, 1 < *n* < 30, and *h* is a significance threshold between 0 and 1. The threshold is set to value *h* = 0.7 for illustration purposes in Figure 2C. For analysis, the threshold value was set to *h* = 0.3 in order to include all perisaccadic STUs that might play a role in transsaccadic integration.

As shown in Figure 2A-C, the modulated subset of STUs (Figure 2B), representing the STUs which are significantly different between the fixation and perisaccadic periods, play a major role in maintaining visuospatial integrity across the saccade. As shown in Figure 2C, replacing the weights of the modulated STUs in the model with their fixation values results in a gap in the readout of the neural responses— and interruption in the detectability and discriminability at RF1 or RF2 in the perisaccadic period.

In the next step, a subset of modulated STUs is identified as contributing to this visuospatial integration—e.g., the continuity of transitioning sensitivity from RF1 to RF2 across the saccade (termed ‘integration-relevant STUs’). The contribution of each modulated STU to the transsaccadic integrity is quantified by evaluating its role in maintaining the sensitivity of the neuron to either the RF1 or RF2 location across a saccadic eye movement, by removing each modulated STU one at a time and testing whether the neuron’s sensitivity decreases. The stimulus kernels of the fitted models (*K_x,y_*(*t,τ*), Figure S3G) reflect changes in the neurons’ spatiotemporal sensitivity at each probe location (*x,y*) and delay (*τ*) across different times to the saccade (*t*). The average spatial sensitivity of the neuron to the stimulus in RF1, *h*_1_(*t*), is quantified as:

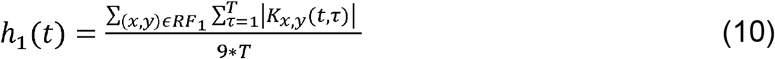

where 1 < *τ* < *T* is the delay parameter in the kernels (in this study *T* = 200 ms as length of stimulus kernels), regarding the history of stimulus from time *t*, and (*x,y*) are the 9 probe locations around center of RF1. The spatial sensitivity index of RF1, *h*_1_(*t*), representing the average sensitivity in terms of the average absolute kernel values, drops after a saccade, while the spatial sensitivity index *h*_2_(*t*) for RF2 increases. A shared sensitivity index (*δ*) across RF1 and RF2 is then defined as the minimum sensitivity to either location, measured across time relative to the saccade. The shared sensitivity is defined as 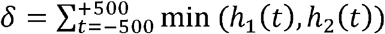 and each modulated STU is considered integration-relevant if nulling its weight results in a decrease in the shared sensitivity of the neuron to the RF1 or RF2.

### Calculating the duration of overlap between RF1 and RF2 discriminability

Figure S6 illustrates how the discriminability at the RF1 and RF2 locations overlap during the perisaccadic period, and how the duration of this overlap time is quantified. The discriminability of the neuron to either RF1 or RF2 (*γ*_1_ and *γ*_2_) is defined as the maximum discriminability across different delays at each response time. The overlap time is then measured as the time where stimuli in both RF1 and RF2 are discriminable. Times *t*_1_ and *t*_2_ representing the start and end of the overlap time are quantified based on the cumulative sum of the baseline subtracted discriminability traces 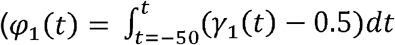 for RF1 and similarly *φ*_2_(*t*) for RF2). Each cumulative function *φ*_1_(*t*) and *φ*_2_(*t*) is calculated for −50 to +200 ms relative to the saccade and normalized to their sum over this period. Time *t*_1_, marking the start of the overlap time, is then measured as the time relative to the saccade when the normalized *φ*(*t*) passes 70% of its maximum. Similarly, time *t*_2_, marking the end of the overlap time, is measured as the time relative to the saccade when the normalized *φ*_2_(*t*) reaches 30% of its maximum. The overlap time is the difference between these start and end times. After removing the integration-relevant STUs the mean population overlap time decreases, and in fact becomes negative, indicating that discriminability around RF1 disappears before discriminability around RF2 arises: a gap in discriminability.

### Future field remapping

It has been shown that in several sensory areas of the cortex, neurons become responsive to their future field (FF, same as RF2) prior to a saccade (in LIP (Colby and Goldberg, 1992), FEF (Sommer and Wurtz, 2006), V2, V3a (Nakamura and Colby, 2002), and V4 (Neupane et al., 2016)). In area MT, predictive future field remapping of stable stimuli has not been observed(Ong and Bisley, 2011), but memory remapping of brief FF stimuli has been reported in MT (Yao et al., 2016) and MST (Inaba and Kawano, 2016). Our results are consistent with the memory-remapping effects previously reported in MT and V4, with neurons responding to brief probes presented in their FF prior to saccade onset. Figure S7A illustrates the population neural response to the visual probes presented in the FF of the neurons in the perisaccadic period (−15 to 0 ms to the saccade onset for the MT neurons and −40 to −10 ms to the saccade onset for the V4 neurons) compared to the probe presentations during fixation (fix1 and fix2, 500 to 200 ms before or after saccade onset respectively), for MT (n=332) and V4 neurons (n= 291). Our results show that the neurons in both areas V4 and MT become sensitive to probes which appear in their future field close to the onset of the saccade. To quantify the FF remapping across the population of the neurons, the future field modulation index (FMI) is defined as:

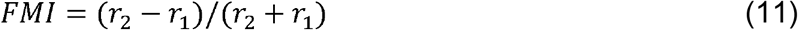

where *r*_2_ is the average firing rate in the late response window (70 to 110 ms to the probe onset for the MT neurons and 65 to 130 ms for the V4 neurons) for the probes appearing in the perisaccadic period and *r*_1_ is the average firing rate of the neurons to the probes presented at the same location on the screen during the first fixation averaged over the same response window. For both the V4 and MT neuronal populations, the modulation index is significantly greater than zero (FMI MT = 0.02±0.01, p=0.001; FMI V4 = 0.06±0.01, p<0.001).

It is important to rule out residual luminance of the FF probe after the eyes have landed as a cause of the response to the perisaccadic FF probes. This “phosphor persistence” effect was determined to be the basis for some previously reported remapping effects (Jonides et al., 1982, 1983). In our case, the sustained residual luminance from a pre-saccadic FF probe would need to last through the duration of the saccade execution to explain the observed responses; this is unlikely given the luminance offset time course of the screen measured with the photodiode (Figure S1). Nevertheless, to provide additional assurance that residual luminance is not responsible for the observed FF remapping, in a different set of experiments, V4 neurons were tested with both white probes on a black background and black probes on a white background (80% contrast). As shown in Figure S7C, for 107 V4 neurons both black and white probes show an FF modulation index significantly greater than zero (white probes: 0.1±0.01, p<0.001, black probes: 0.06±0.01, p=0.001), demonstrating that residual luminance does not account for the FF remapping effect. The FMI is calculated for the black probes presented −20 to +10 ms relative to the saccade onset and in the response window of 65 to 105 ms relative to the saccade onset. The FF modulation index for white and black probes are positively correlated across neurons (Pearson correlation, r = 0.31, p<0.001), consistent with remapping based on spatial rather than luminance properties of the stimulus. Thus, both the photodiode-measured time-course of stimulus offset and the presence of remapping for black probes on a white background indicate that phosphor persistence is not responsible for the observed FF remapping phenomenon.

We also considered the possibility of a role of persistent phosphor luminance in generating the RF persistence effect. This possibility is slim since the RF persistence is evoked by the presence of a probe in the RF1 right before the saccade, so even if there was phosphor persistence it would be removed from the RF of the neuron by the subsequent saccade. However, as an additional control we confirmed the presence of the RF persistence for the black probes presented on the white background (PMI_V4_ = 0.05±0.02, p=0.001 for 107 V4 neurons).

### Timing of the perisaccadic modulation in areas MT and V4

Figure S8A-B compares the response map for a sample V4 (Figure S8A) and MT (Figure S8B) neuron in the early response window (V4 50 to 60 ms and MT 50 to 70 ms relative to the probe onset) versus the late window (V4 60 to 130 ms and MT 80 to 90 ms relative to the probe onset) in the fixation (500 to 200 ms before the saccade onset) and perisaccadic (−10 to +3 ms relative to the saccade onset) periods. The response map in the late window of the perisaccadic period shows the RF persistence effect in both areas. To calculate the significance of the perisaccadic modulation of RF1 responses over different response windows, the firing rates in response to the RF1 probe in the perisaccadic period were compared to those in the fixation period using a t-test. Figure S8 plots the significance of this comparison (t-score) for a sliding 30 ms response window relative to the probe onset. Positive t-score values correspond to an enhancement in the perisaccadic firing rate and the negative values indicate a suppression in the perisaccadic firing rates.

### Perisaccadic modulation of response in congruent versus incongruent saccades

As shown in Figure 4E, the PMI is significantly greater than zero for the neurons in area MT regardless of the direction of the saccade. The congruent saccades are defined as those in which saccade-induced retinal motion is within 180 degrees of the neuron’s preferred motion direction (incongruent saccades induce motion >180 degrees from the neuron’s preferred direction of motion). In Figure 4E, we show that MT neurons’ perisaccadic responses are modulated for saccades that are either congruent or incongruent with the neuron’s preferred motion direction. As shown in Figure S9, for 61 MT neurons recorded during saccades in opposite directions, the PMI values for both congruent and incongruent saccades are significantly greater than zero (congruent PMI=0.04±0.02, p=0.024, incongruent PMI=0.07±0.02, p=0.005), and the PMI is not different between two opposite directions of saccade (congruent versus incongruent, p=0.273).

### Saccade target remapping

Whereas the early component of probe-aligned response mostly reflects the classic RF of the neurons, we found that around the time of saccade the late component of response shows sensitivity to other parts of visual space. As shown in Figure S7, some neurons become sensitive to RF2 probes in the perisaccadic period (FF remapping). Some neurons show RF persistence (Figure 4), a late component of the response to perisaccadic RF1 probes. A late component of the response is also observed for perisaccadic probes around the ST (ST remapping). The ST remapping, FF remapping, and RF persistence phenomena are observed in varying combinations across the neuronal population. Figure S10A shows the response pattern for 3 sample neurons which become sensitive to ST probes in the perisaccadic period, and compares it to the pre- and post-saccadic RFs. The late component of the response is shown for the perisaccadic period in Figure S10A, illustrating how ST remapping occurs at the same time as FF remapping or RF persistence. The pre-(left) and post-(middle) saccadic RFs of the neuron are calculated by averaging the response to probes presented more than 100 ms before or after the saccade onset across a 30 ms window around the maximum response time relative to the probe onset. For the perisaccadic maps the firing rate of the neurons are averaged for the probes presented −10 to +3, −10 to −3, and −10 to −3 ms relative to saccade onset and the response windows are 105-115, 65-90, and 80-90 ms relative to probe onset.

Figure S10B-D shows the detectability map for ST probes across the time to saccade for a sample neuron (Figure S10B), the population of neurons (Figure S10C), and versus the RF1 and RF2 detectability (Figure S10D). Figure S10B shows the detectability map for a sample ST probe for a sample neuron with ST remapping. To measure the population detectability maps in Figure S10C-D, we first identify the probe locations near the saccade target that do not overlap with RF1 or RF2, or the vector connecting them. More specifically, first we select all the probes around the saccade target if they are within a window of 50% of the saccade vector centered at the ST of each neuron. Then, to avoid any confound with the conventional RF or the corresponding retinal locations during the eye movement, probes are discarded if they are closer than 2 dva to any point along the vector connecting the RF1 probe to the RF2. Next we determined which of these probes near the ST are evoking significantly different responses during the perisaccadic vs. fixation periods. We first used an ANOVA test on the difference of the firing rate of the neurons in the late window of the perisaccadic versus average of the fixation period before the saccade in response to all ST probes. If the responses across all the ST probes are modulated perisaccadically for a given neuron (p<0.01, ANOVA), we then use a post hoc Bonferroni test to identify the modulated probes (p<0.05). Plots in Figure S10 C and D are averaged across modulated probes across the population of 51 V4 and MT neurons. The ST discriminability contour shows where the detectability map crosses threshold of 0.55. Figure S10D shows detectability at the RF1, RF2, and ST locations, across the time of probe and response relative to the saccade.

Figure S10E shows the co-occurrence of the RF persistence, FF remapping, and ST remapping phenomena in the population of neurons. A neuron is considered to be modulated if the firing rates evoked by RF, FF, or ST probes in the perisaccadic period are significantly different from firing rates during the fixation period, using a Wilcoxon rank-sum test with significance level of 0.05. RF, FF, and ST effects are defined as described in the main text and previous sections of STAR Methods. For the ST remapping effect, neurons were only included in the analysis if the ST probes were far from the neuron’s FF, to avoid potentially confounding the FF response with an ST modulation. To show that the FF modulation also exists independent of the ST modulation, we measured the FMI for the neurons where the FF of the neuron is far from the ST (distance between FF probe and ST > 8 dva). The results show that the FF remapping exists for these neurons in both MT and V4 areas (FMI_V4_ = 0.06±0.01, p=0.001, n=93, FMI_MT_ =0.06±0.01, p<0.001, n=216) showing the coexistence of modulations around both the ST and FF.

## QUANTIFICATION AND STATISTICAL ANALYSIS

All quantification and statistical analysis were performed with custom MATLAB scripts. All reported p-values are from a Wilcoxon signed-rank test unless otherwise stated. All statistics are reported as mean ± SE (standard error) unless otherwise stated. The exact number of samples used in the statistical test is mentioned at the instance that it has been used in the text.

## DATA AND CODE AVAILABILITY

All neuronal data and the computational source codes are available for review process at http://gofile.me/2d2XM/u3G5eCltx. The data will be available for public use after acceptance.

## SUPPLEMENTAL INFORMATION

**Figure S1, related to Figure 1.** Details of visual stimuli and RFs. (A) Spatial extent and positioning of the probe grids across different sessions (black squares). Black dots represent the saccade target positions and the gray dot marks the FP. (B) Mean probe luminance over time aligned to probe offset (measured by a photodiode averaged across 810 trials of a sample session, error bars not visible). (C) RF center locations for the population of V4 (left) and MT neurons (right); colors indicate the animal.

**Figure S2, related to Figure 1.** Dimensionality reduction procedure for STU selection. (A) Temporal distribution of STUs with a significant stimulus-response relationship (gray) for a probe inside RF1 of a sample neuron. The STUs for the RF1 probe occur primarily at delays around the latency of the sample neuron (50-70 ms, x-axis) and for probes appearing before saccade onset (−540 to 0 ms, y-axis). After the saccade, prevalence of the STUs for RF1 probes diminishes, showing that the eye movement has moved this probe out of neuron’s RF. The distribution of fitted weights for different subsets of trials is shown for sample STUs with non-significant (orange) and significant (green) stimulus-response relationships. The histograms show the distribution of fitted weights of a simplified single STU LNP model across different random subsets of trials (n = 100 random selections) versus the fitted control weights where trials are randomly shuffled. (B) Mean fitted weights for each STU of RF1 of the sample neuron in A, fitted to the data (x-axis) and shuffled (y-axis; mean of Y random subsets). STU locations with non-significant stimulus response relationships are shown in orange while the rest of the STUs are shown in green.

**Figure S3, related to Figure 1.** Schematic of sparse generalized linear model structure and fitted components. (A) The input to the model is shown by a set of stimulus sequences along each spatial dimension for 3 sample locations. (B) S-kernels representing the neuron’s sensitivity at different times. (C) The offset kernel representing the changes in the neuron’s baseline activity across time relative to the saccade. (D) Post-spike kernel representing neuron’s inherent features like refractoriness or burstiness. (E) The nonlinear function. The input to the model gets convolved with the S-kernels and is then summed with the offset kernel and the feedback signal through the post-spike kernel. The resulting signal is then passed through the sigmoidal nonlinearity to generate a predicted instantaneous firing rate of the neuron. (F) The predicted firing rate is then passed through a conditionally Poisson generator to generate the predicted spike train. (G) The fitted S-kernel for a sample neuron for RF1 (top) and RF2 (bottom) locations. (H) The fitted offset kernel (left) and post-spike kernel (right) for the sample neuron. The gray traces show the fitted kernel to the randomly chosen subsets of the data (65% of the data).

**Figure S4, related to Figure 1**. Performance of the model in predicting the neural responses. (A) Stimulus aligned neural response (black) and the prediction of the model (red) for a sample MT neuron. The response of the neuron is averaged over repetitions of the probe presentations inside the neuron’s RF in the fixation period (500 to 200 ms before the saccade onset) and smoothed with a Gaussian window of 20 ms FWHM. The error bars are calculated over repetitions of the probe across trials. (B) The percentage of the explained variance (EV) for the model-predicted firing rate for all neurons in area MT and V4 (n = 623), for the stimulus-aligned neural response when a stimulus is presented inside the RF during the fixation period. In B,C and E the red triangles indicate medians; histograms show the marginal distributions (left, bottom) and the difference (upper-right). (C) The normalized cross-correlation (CC) between the model-predicted response and the data, compared to the data-data correlation, averaged over different sequences of stimuli in all conditions for the population of 623 MT and V4 neurons. (D) The traces show the model-predicted firing rate (dashed line) versus the recorded neural response (solid line) for sample trials which were of good fit (left; best LL) and average fit (right, median LL) for two sample neurons (top and bottom). Spikes in each trial shown as black vertical lines below the smoothed traces. (E) Comparison of the LL of the model predicted firing rate and the recorded neural response in the perisaccadic versus fixation period.

**Figure S5, related to Figure 2.** Detectability across delay and time to saccade. (A) Illustration of the method for calculating the detectability of a sample probe *s*_52_ at a specific time to saccade (*t*\*) and delay relative to probe presentation (*τ*\*). Top: White squares show the location of three example visual probes, for probes *s*_27_, *s*_52_, *s*_12_ where *s*_1_,…*s*_81_ represent the 81 possible locations on the 9 by 9 stimulus grid. Middle: Example stimulus sequences, in which either sample probe *s*_52_ appears at a specific delay relative to *t\** (top three rows), or a random probe appears at the same delay relative to *t*\* (bottom three rows). Bottom: Histograms show the firing rate distributions for the trials in which sample probe *s*_52_ appears at a specific delay relative to *t*\* (red), or a random probe appears at the same delay (blue). The AUC of an ROC between these two distributions for each time and delay value produces the detectability values shown in B-D. (B) Detectability of an example MT neuron across delay values for one time to saccade (*t*\* = +10 ms). Detectability peaks at the response latency of the neuron (*τ*\* = 60 ms). (C) Detectability for the same example neuron shown in B, across both delay and time to saccade. Horizontal dotted line corresponds to the *t*\* plotted in B; vertical dotted line corresponds to delay with maximum detectability. (D) Detectability for the same neuron shown in B and (C) plotted as a function of time between stimulus and saccade (y-axis) and response relative to saccade (x-axis), for probe locations in RF1 (left) and RF2 (right). Outlines show the period when the AUC > 0.61. The projected lines on top show the times that the detectability threshold was exceeded for RF1 (blue) and RF2 (red). (E) Average detectability map over the population of the neurons measured based on the recorded neural responses. (F) Average discriminability map over the population of neurons measured based on the recorded neural responses.

**Figure S6, related to Figure 2.** Identifying the overlap time with and without the modulated STUs and determining the integration-relevant STUs. (A) Discriminability metric of a sample neuron around RF1 (blue) and RF2 (red) over time relative to the saccade. Time *t*_1_ (vertical dashed line) and *t*_2_ (vertical solid line) represent the start and the end of the overlap time. (B) Overlap time across the population of the neurons with and without the integration-relevant STUs. The red triangles and the dashed lines indicate medians; histograms show the marginal distributions (left, bottom) and the difference (upper-right). (C) Sensitivity index of the neuron to probes around RF1 (blue) and RF2 (red) over time relative to the saccade. The sensitivity of the neuron to the RF1 drops after the saccade while the sensitivity to the probes around RF2 increases. The combined sensitivity, measured as the shared area under the curve of both RF1 and RF2 sensitivity, is shown in gray. If eliminating an STU resulted in a decrease in the combined sensitivity, it was deemed integration-relevant. (D) Effect of eliminating integration-relevant STUs on the discriminability (left) and detectability (right) for the neuronal population. Contour lines show time periods in which discriminability (left) and detectability (right) at the RF1 (blue) and RF2 (red) locations exceeded the same threshold used in Figure 2. Projections on top show a gap in discriminability/detectability during the perisaccadic period.

**Figure S7, related to Figure 3.** Future field remapping in areas V4 and MT. (A) The probe aligned response in area V4 (left) and MT (right), for the white probes presented in RF2 during the pre-, peri-, and post-saccadic periods (blue, black, and red respectively). The dashed lines indicate the response window used for the FMI calculation. (B) The probe aligned response in area V4 for the black probes presented in RF2 during the pre-, peri-, and post-saccadic periods (blue, black, and red respectively). (C) FMI for the black probes on the white background versus the white probes on the black background over 107 V4 neurons.

**Figure S8, related to Figure 4.** Spatial map of the perisaccadic late response in V4 and MT neurons and robustness of the perisaccadic modulation across different delays. Response map of a sample (A) V4 and (B) MT neuron is shown in early (left) and late (right) response windows for fixation (top) and perisaccadic (bottom) periods. (C) Plot shows the timing of the perisaccadic modulation (V4: blue and MT: red) measured as the t-score of the average firing rate in response to the RF1 probe in the perisaccadic period versus the fixation period, for different response windows relative to the probe onset. The x-axis value indicates the start of the response window relative to the probe onset (window duration 30 ms).

**Figure S9, related to Figure 4.** Saccade direction does not alter the strength of the late response to perisaccadic RF1 stimuli in MT. The scatter plot shows the PMI for the saccades congruent versus incongruent with the preferred direction of 61 MT neurons. For each data point in the scatter plot, the response of a single neuron is considered for saccades in two opposite directions. The histograms along the x and y axis show the distribution of PMI values for congruent and incongruent saccades. The upper right histogram shows the difference of the PMI for congruent versus incongruent saccades.

**Figure S10, related to Figure 4.** Coexistence of RF, FF, and ST modulations. (A) Three sample neurons responding to perisaccadic probes around the ST location. Heat maps show the response to probes at different locations before (left), after (middle), and around (right) the saccade onset. The fixation point (black) and saccade target (red) are shown as filled circles. (B) Detectability of the sample neuron for a probe around the ST. (C) Detectability of the population for the probes around the ST for the population of 51 neurons. The contour is showing where the detectability exceeds a threshold. (D) The detectability of the RF1, RF2, and ST probes for the population. The projection on top is showing where the detectability exceeds a threshold for RF1, RF2, and ST. (E) Venn diagram showing the number of neurons exhibiting perisaccadic modulation at the RF, FF, and ST probe locations. 333 out of 623 neurons show modulation at one or more of these locations.

