## Supplemental Information for "A Sensory Memory to Preserve Visual Representations Across Eye Movements"

Supplemental Information consists of:

Supplemental figures S1-S10

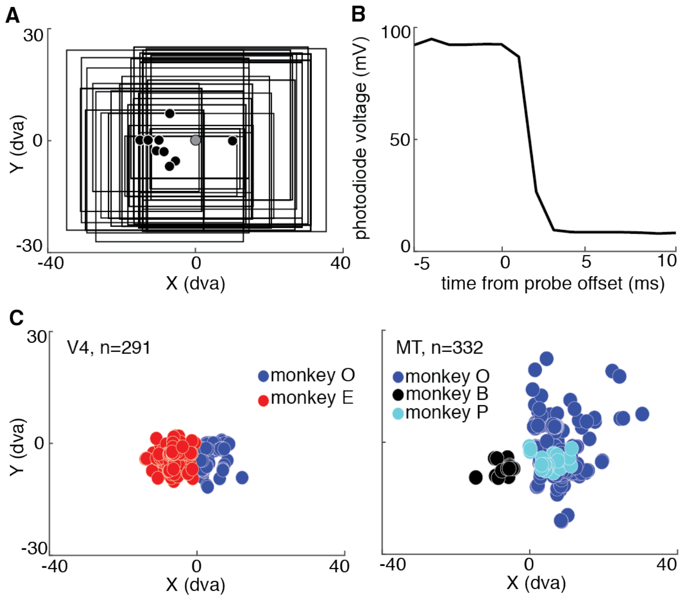


**Figure S1, related to Figure 1.** Details of visual stimuli and RFs. (A) Spatial extent and positioning of the probe grids across different sessions (black squares). Black dots represent the saccade target positions and the gray dot marks the FP. (B) Mean probe luminance over time aligned to probe offset (measured by a photodiode averaged across 810 trials of a sample session, error bars not visible). (C) RF center locations for the population of V4 (left) and MT neurons (right); colors indicate the animal.

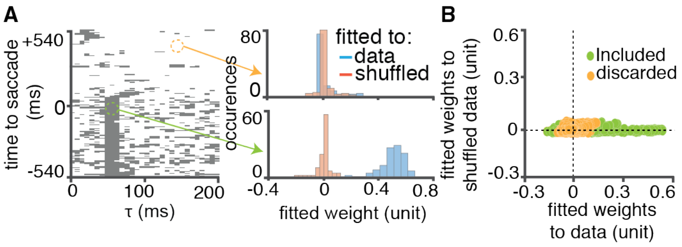


**Figure S2, related to Figure 1.** Dimensionality reduction procedure for STU selection. (A) Temporal distribution of STUs with a significant stimulus-response relationship (gray) for a probe inside RF1 of a sample neuron. The STUs for the RF1 probe occur primarily at delays around the latency of the sample neuron (50-70 ms, x-axis) and for probes appearing before saccade onset (-540 to 0 ms, y-axis). After the saccade, prevalence of the STUs for RF1 probes diminishes, showing that the eye movement has moved this probe out of neuron's RF. The distribution of fitted weights for different subsets of trials is shown for sample STUs with non-significant (orange) and significant (green) stimulus-response relationships. The histograms show the distribution of fitted weights of a simplified single STU LNP model across different random subsets of trials (n = 100 random selections) versus the fitted control weights where trials are randomly shuffled. (B) Mean fitted weights for each STU of RF1 of the sample neuron in A, fitted to the data (x-axis) and shuffled (y-axis; mean of Y random subsets). STU locations with non-significant stimulus response relationships are shown in orange while the rest of the STUs are shown in green.

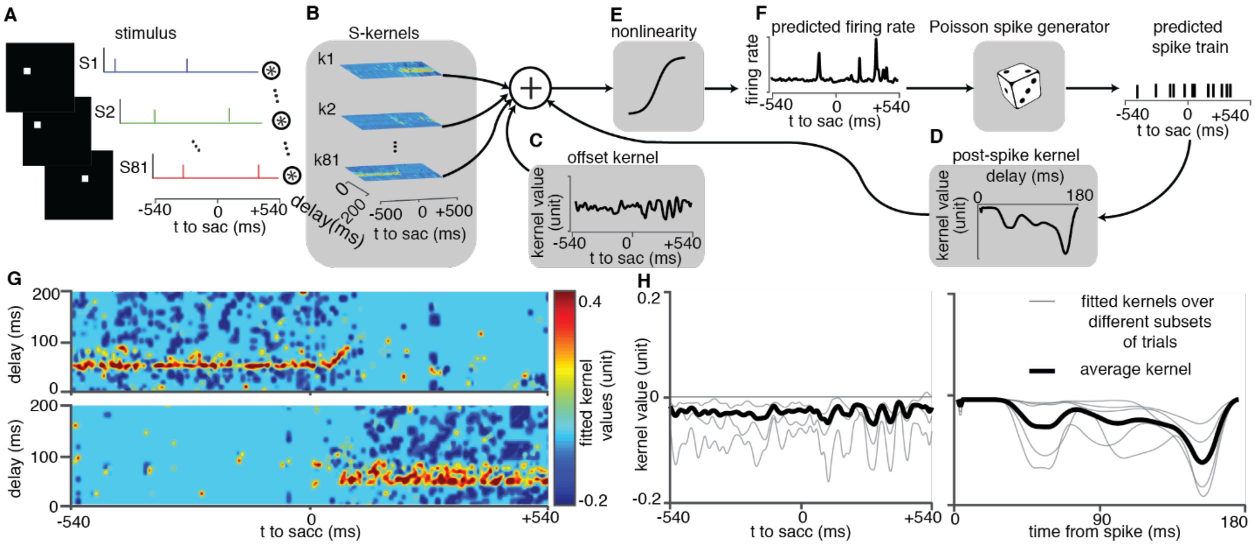


**Figure S3, related to Figure 1.** Schematic of sparse generalized linear model structure and fitted components. (A) The input to the model is shown by a set of stimulus sequences along each spatial dimension for 3 sample locations. (B) S-kernels representing the neuron's sensitivity at different times. (C) The offset kernel representing the changes in the neuron's baseline activity across time relative to the saccade. (D) Post-spike kernel representing neuron's inherent features like refractoriness or burstiness. (E) The nonlinear function. The input to the model gets convolved with the S-kernels and is then summed with the offset kernel and the feedback signal through the post-spike kernel. The resulting signal is then passed through the sigmoidal nonlinearity to generate a predicted instantaneous firing rate of the neuron. (F) The predicted firing rate is then passed through a conditionally Poisson generator to generate the predicted spike train. (G) The fitted S-kernel for a sample neuron for RF1 (top) and RF2 (bottom) locations. (H) The fitted offset kernel (left) and post-spike kernel (right) for the sample neuron. The gray traces show the fitted kernel to the randomly chosen subsets of the data (65% of the data).

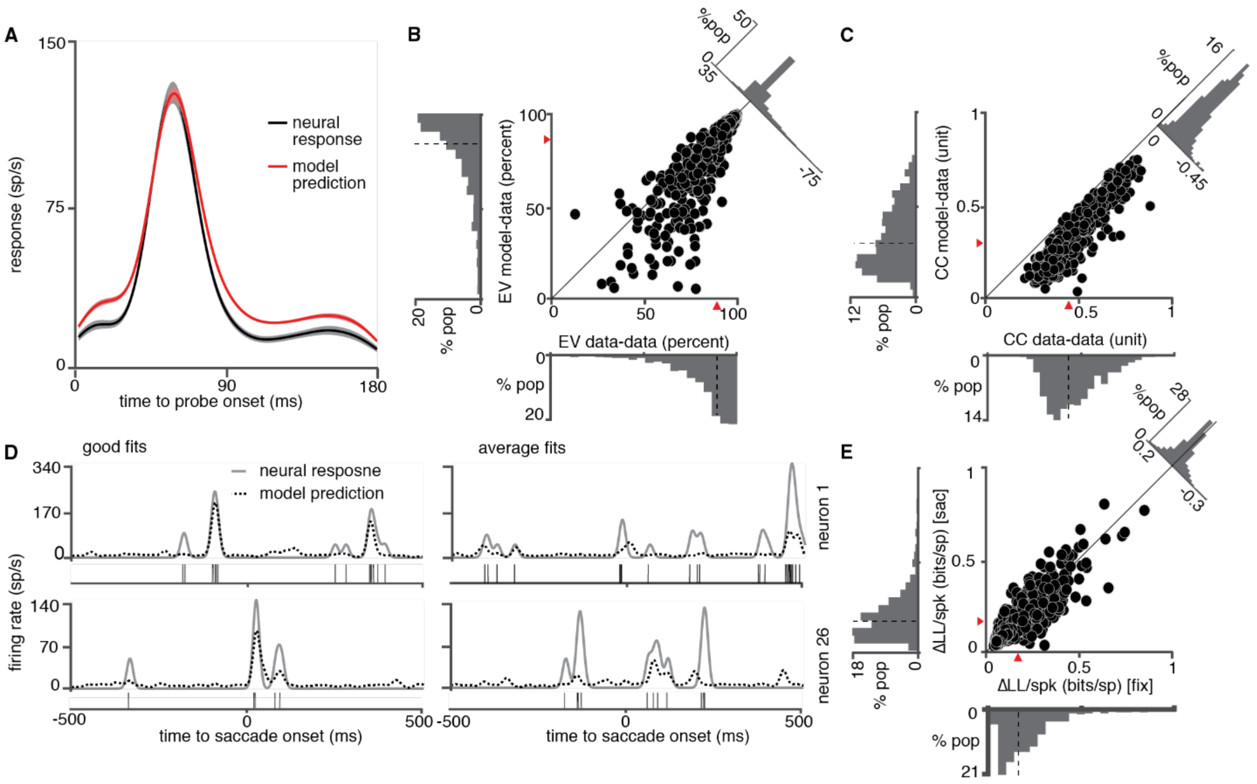


**Figure S4, related to Figure 1**. Performance of the model in predicting the neural responses. (A) Stimulus aligned neural response (black) and the prediction of the model (red) for a sample MT neuron. The response of the neuron is averaged over repetitions of the probe presentations inside the neuron's RF in the fixation period (500 to 200 ms before the saccade onset) and smoothed with a Gaussian window of 20 ms FWHM. The error bars are calculated over repetitions of the probe across trials. (B) The percentage of the explained variance (EV) for the model-predicted firing rate for all neurons in area MT and V4 (n = 623), for the stimulus-aligned neural response when a stimulus is presented inside the RF during the fixation period. In B,C and E the red triangles indicate medians; histograms show the marginal distributions (left, bottom) and the difference (upper-right). (C) The normalized cross-correlation (CC) between the model-predicted response and the data, compared to the data-data correlation, averaged over different sequences of stimuli in all conditions for the population of 623 MT and V4 neurons. (D) The traces show the model-predicted firing rate (dashed line) versus the recorded neural response (solid line) for sample trials which were of good fit (left; best LL) and average fit (right, median LL) for two sample neurons (top and bottom). Spikes in each trial shown as black vertical lines below the smoothed traces. (E) Comparison of the LL of the model predicted firing rate and the recorded neural response in the perisaccadic versus fixation period.

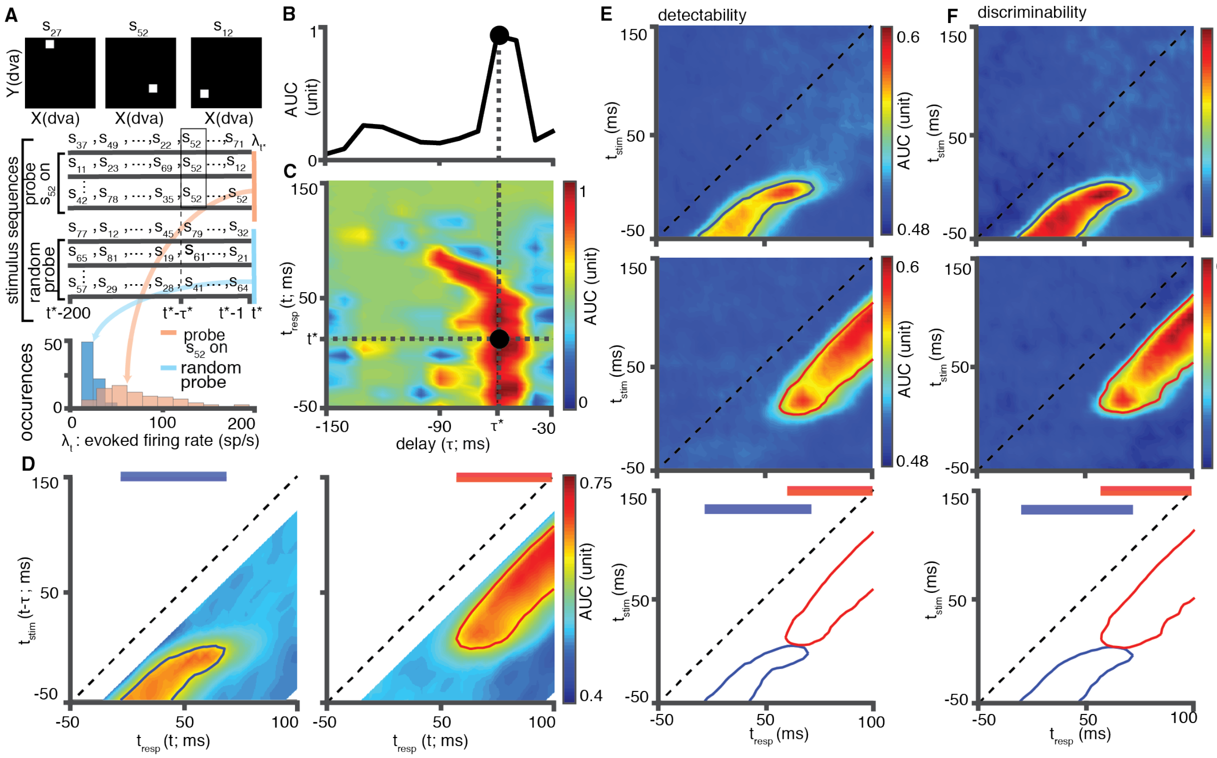


**Figure S5, related to Figure 2.** Detectability across delay and time to saccade. (A) Illustration of the method for calculating the detectability of a sample probe $s_{52}$ at a specific time to saccade ($t^*$) and delay relative to probe presentation ($\tau^*$). Top: White squares show the location of three example visual probes, for probes $s_{27}$ , $s_{52}$ , $s_{12}$ where $s_1, \dots s_{81}$ represent the 81 possible locations on the 9 by 9 stimulus grid. Middle: Example stimulus sequences, in which either sample probe $s_{52}$ appears at a specific delay relative to *t** (top three rows), or a random probe appears at the same delay relative to *t** (bottom three rows). Bottom: Histograms show the firing rate distributions for the trials in which sample probe $s_{52}$ appears at a specific delay relative to *t** (red), or a random probe appears at the same delay (blue). The AUC of an ROC between these two distributions for each time and delay value produces the detectability values shown in B-D. (B) Detectability of an example MT neuron across delay values for one time to saccade (*t** =

+10 ms). Detectability peaks at the response latency of the neuron ($\tau^*$ = 60 ms). (C) Detectability for the same example neuron shown in B, across both delay and time to saccade. Horizontal dotted line corresponds to the $t$* plotted in B; vertical dotted line corresponds to delay with maximum detectability. (D) Detectability for the same neuron shown in B and (C) plotted as a function of time between stimulus and saccade (y-axis) and response relative to saccade (x-axis), for probe locations in RF1 (left) and RF2 (right). Outlines show the period when the AUC > 0.61. The projected lines on top show the times that the detectability threshold was exceeded for RF1 (blue) and RF2 (red). (E) Average detectability map over the population of the neurons measured based on the recorded neural responses. (F) Average discriminability map over the population of neurons measured based on the recorded neural responses.

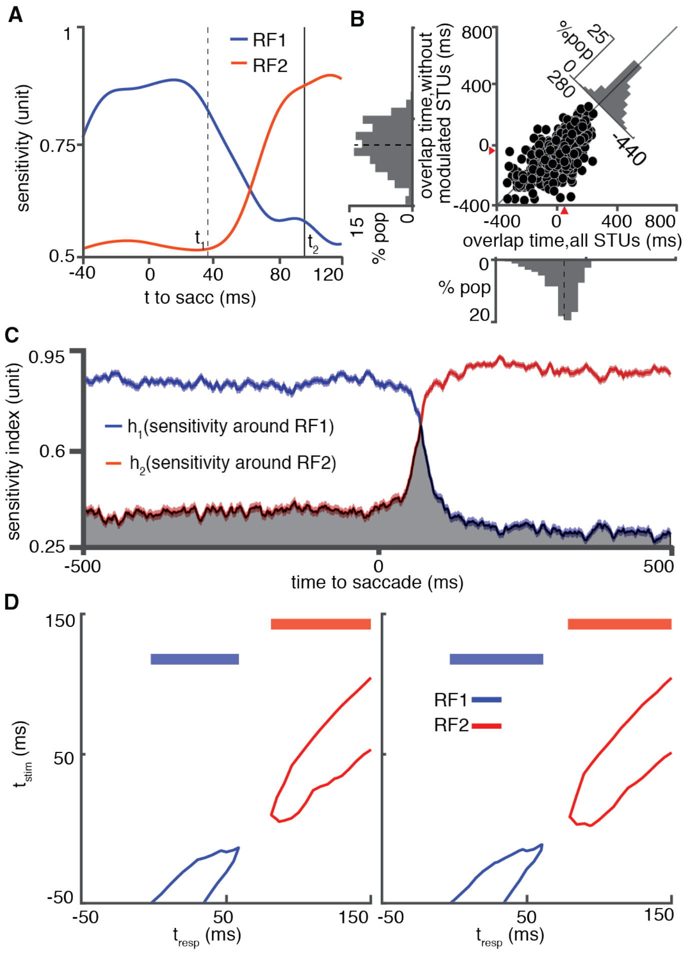


**Figure S6, related to Figure 2.** Identifying the overlap time with and without the modulated STUs and determining the integration-relevant STUs. (A) Discriminability metric of a sample neuron around RF1 (blue) and RF2 (red) over time relative to the saccade. Time $t_1$ (vertical dashed line) and $t_2$ (vertical solid line) represent the start and the end of the overlap time. (B) Overlap time across the population of the neurons with and without the integration-relevant STUs. The red triangles and the dashed lines indicate medians; histograms show the marginal distributions (left, bottom) and the difference (upper-right). (C) Sensitivity index of the neuron to probes around RF1 (blue) and RF2 (red) over time relative to the saccade. The sensitivity of the neuron to the RF1 drops after the saccade while the sensitivity to the probes around RF2 increases. The combined sensitivity, measured as the shared area under the curve of both RF1 and RF2 sensitivity, is shown in gray. If eliminating an STU resulted in a decrease in the combined sensitivity, it was deemed integration-relevant. (D) Effect of eliminating integration-relevant STUs on the discriminability (left) and detectability (right) for the neuronal population. Contour lines show time periods in which discriminability (left) and detectability (right) at the RF1 (blue) and RF2 (red) locations exceeded the same threshold used in figure 2. Projections on top show a gap in discriminability/detectability during the perisaccadic period.

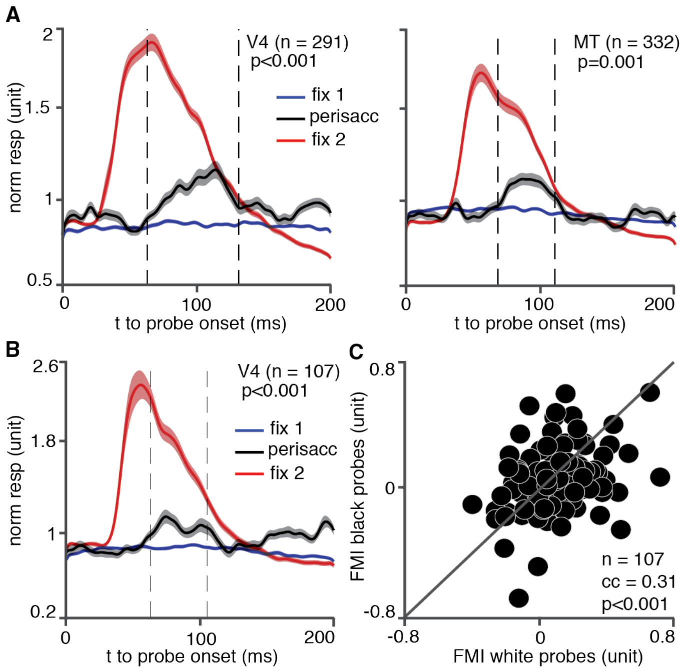


**Figure S7, related to Figure 3.** Future field remapping in areas V4 and MT. (A) The probe aligned response in area V4 (left) and MT (right), for the white probes presented in RF2 during the pre-, peri-, and post-saccadic periods (blue, black, and red respectively). The dashed lines indicate the response window used for the FMI calculation. (B) The probe aligned response in area V4 for the black probes presented in RF2 during the pre-, peri-, and post-saccadic periods (blue, black, and red respectively). (C) FMI for the black probes on the white background versus the white probes on the black background over 107 V4 neurons.

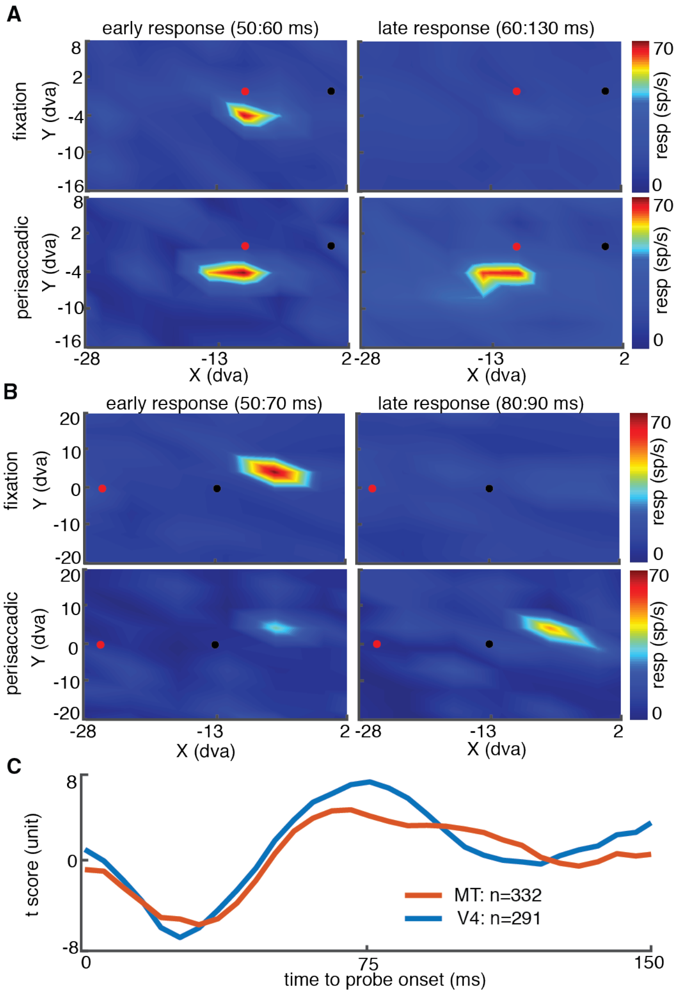


**Figure S8, related to Figure 4.** Spatial map of the perisaccadic late response in V4 and MT neurons and robustness of the perisaccadic modulation across different delays. Response map of a sample (A) V4 and (B) MT neuron is shown in early (left) and late (right) response windows for fixation (top) and perisaccadic (bottom) periods. (C) Plot shows the timing of the perisaccadic modulation (V4: blue and MT: red) measured as the t-score of the average firing rate in response to the RF1 probe in the perisaccadic period versus the fixation period, for different response windows relative to the probe onset. The x-axis value indicates the start of the response window relative to the probe onset (window duration 30 ms).

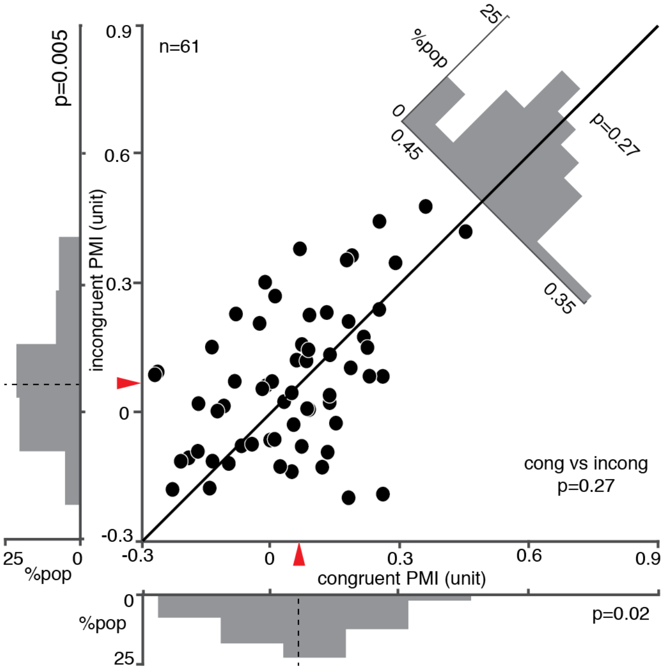


**Figure S9, related to Figure 4.** Saccade direction does not alter the strength of the late response to perisaccadic RF1 stimuli in MT. The scatter plot shows the PMI for the saccades congruent versus incongruent with the preferred direction of 61 MT neurons. For each data point in the scatter plot, the response of a single neuron is considered for saccades in two opposite directions. The histograms along the x and y axis show the distribution of PMI values for congruent and incongruent saccades. The upper right histogram shows the difference of the PMI for congruent versus incongruent saccades.

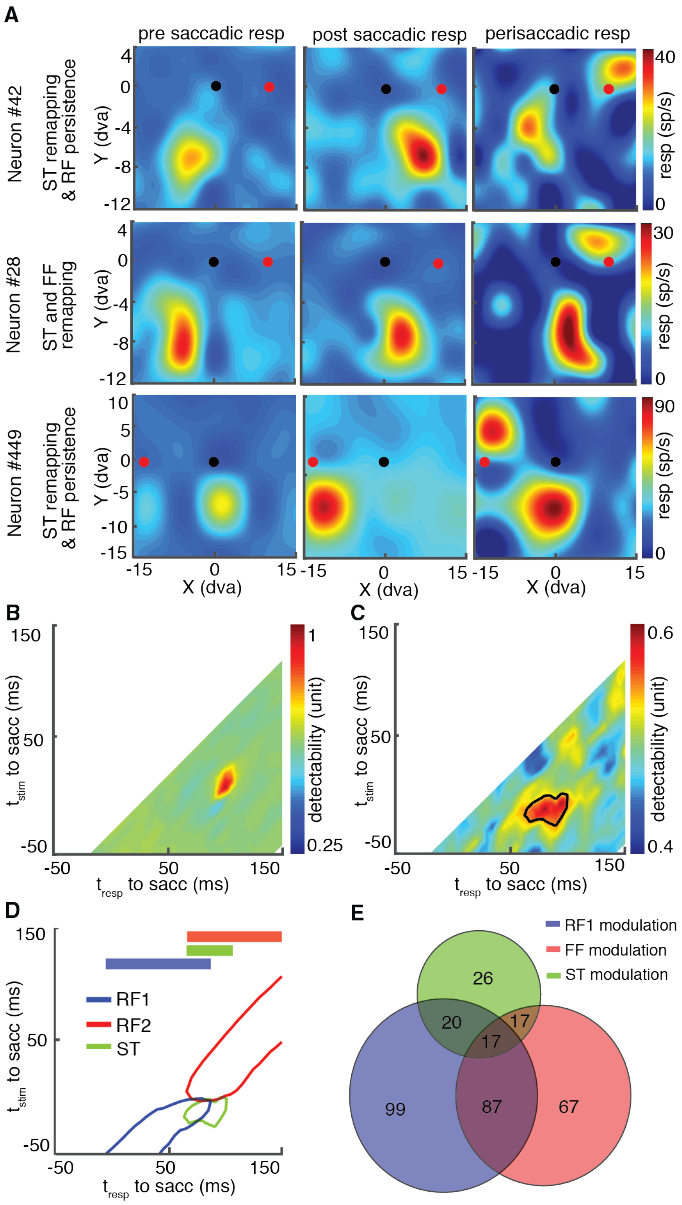


**Figure S10, related to Figure 4.** Coexistence of RF, FF, and ST modulations. (A) Three sample neurons responding to perisaccadic probes around the ST location. Heat maps show the response to probes at different locations before (left), after (middle), and around (right) the saccade onset. The fixation point (black) and saccade target (red) are shown as filled circles. (B) Detectability of the sample neuron for a probe around the ST. (C) Detectability of the population for the probes around the ST for the population of 51 neurons. The contour is showing where the detectability exceeds a threshold. (D) The detectability of the RF1, RF2, and ST probes for the population. The projection on top is showing where the detectability exceeds a threshold for RF1, RF2, and ST. (E) Venn diagram showing the number of neurons exhibiting perisaccadic modulation at the RF, FF, and ST probe locations. 333 out of 623 neurons show modulation at one or more of these locations.
